# Dense CRISPR Mapping of the WAVE Complex Identifies Pharmacological Modulators of AD-Linked Myeloid Migration

**DOI:** 10.64898/2026.06.30.735672

**Authors:** Jason C. Ngo, Yice Xu, Marta Olah, Falak Sher

**Affiliations:** Center for Translational and Computational Neuroimmunology, Columbia University Irving Medical Center, New York, NY USA; Taub Institute for Research on Alzheimer’s Disease and Aging Brain, Columbia University Irving Medical Center, New York, NY USA; Department of Neurology, Columbia University Irving Medical Center, New York, NY USA; Department of Biological Sciences, Graduate School of Arts and Sciences, Columbia University, New York, NY 10027, USA

## Abstract

Although genetic risk for Alzheimer’s disease (AD) strongly converges on microglial pathways, the druggable functional protein regions that control disease-relevant microglial behaviors remain largely unknown. Here, we applied dense CRISPR-Cas9 mutagenesis and CRISPRtile-based functional mapping to the WAVE regulatory complex (WRC), a central regulator of actin remodeling and cell migration. In a pooled CCL2-directed migration assay in human THP-1 myeloid cells, perturbation of *NCKAP1L*, *CYFIP1*, and *BRK1* impaired migration and revealed divergent effects among WRC paralogs. Residue-level analysis mapped discrete migration-associated functional regions within CYFIP1 and NCKAP1L, including a CYFIP1 regulatory hotspot and a prioritized NCKAP1L region nominated for pharmacological targeting. Human single-nucleus datasets identified *NCKAP1L* as a microglia-enriched WRC component expressed across diverse microglial states. Machine-learning-guided compound prediction nominated Montelukast sodium and Piperacetazine, which we experimentally validated as negative and positive modulators of chemokine-directed migration, respectively. These findings establish WRC-dependent migration as a pharmacologically tunable myeloid process relevant to AD.

## Introduction

Alzheimer’s disease (AD) is the leading cause of dementia worldwide, affecting tens of millions of people and imposing an enormous and growing burden on patients, caregivers, and healthcare systems [1, 52]. Extensive investigation of amyloid-β and tau pathology has provided fundamental insights into disease mechanisms and has guided the development of therapeutic strategies targeting these hallmark features of AD [10, 23]. Nevertheless, the incomplete clinical efficacy observed to date and the substantial heterogeneity among patients suggest that additional biological pathways contribute to disease onset and progression [14, 57, 58]. Consequently, increasing attention has focused on identifying complementary disease-modifying mechanisms, particularly those that influence an individual’s genetic susceptibility to AD and shape cellular responses to neurodegenerative pathology [8, 33].

Genome-wide association studies (GWAS) have been central to this shift. Of the dozens of risk loci now associated with AD, a striking proportion lie in or near genes that are highly or selectively expressed in microglia, the resident immune cells of the brain [15, 18, 21, 32, 36]. This pattern suggests that microglial dysfunction is not merely a reactive consequence of amyloid and tau accumulation, but may act as a primary contributor of disease susceptibility and progression. Among the implicated genes, rare coding variants in *TREM2*, *PLCG2*, and *ABI3* directly alter microglial innate immune signaling, providing some of the strongest causal evidence linking microglial cell biology to AD risk [48]. These findings reinforce the premise that the molecular machinery governing microglial behavior, including environmental sensing, chemotaxis, and cellular responsiveness, represents a viable domain for therapeutic intervention.

One such piece of machinery is the WAVE regulatory complex (WRC), a heteropentameric assembly composed of WAVE proteins (*WASF1, WASF2, WASF3), CYFIP1/CYFIP2, NCKAP1/NCKAP1L* (HEM), *ABI1/ABI2/ABI3*, and *BRK1* (HSPC300). The WRC is the principal activator of the Arp2/3 complex, driving branched actin polymerization that underlies cell migration, morphological remodeling, and phagocytic cup formation [17, 39, 40]. Notably, several WRC subunits have independent links to neurological diseases. For instance, ABI3 carries rare coding variants associated with AD risk [48], *CYFIP1* and *CYFIP2* are implicated in autism, schizophrenia, and dementia [38, 50], *NCKAP1* expression is altered in AD brain [55]. Although, WRC subunits are robustly expressed in human microglia, and recent work has begun to define roles for individual components, particularly *CYFIP1* and the Arp2/3 axis [45] in microglial morphology and surveillance behavior. However, whether and how the WRC coordinates directed microglial chemotaxis toward disease-relevant chemoattractants like CCL2, a critical chemokine orchestrating myeloid cell recruitment in AD pathologies, remains essentially unexplored. Consequently, determining if microglial WRC activity can be pharmacologically tuned in a bidirectional manner presents a highly promising yet unexploited avenue for therapeutic development.

Addressing this gap requires resolution beyond what conventional gene-level CRISPR screens can offer. A screen that identifies a WRC subunit as a “hit” cannot, on its own, indicate which residues or structural regions drive the phenotype, nor whether those regions correspond to surfaces that could plausibly be targeted by a small molecule. We have previously developed and applied CRISPR-Cas9 saturating/tiling mutagenesis to address exactly this kind of question: dissecting a minimal functional enhancer within the *BCL11A* locus at near-nucleotide resolution [3], identifying druggable subunit interfaces within a chromatin-modifying complex through comprehensive in situ mutagenesis [46], and building computational frameworks, CRISPR-SURF [26], CRISPRO [43], to extract functional and structural signal from tiling screen data. Building on this foundation, we recently introduced CRISPRtile, an AI-enabled platform that converts dense mutagenesis data into functional and toxicity landscapes and predicts small molecules capable of phenocopying favorable perturbations, which we previously demonstrated by mapping the NLRP3 inflammasome [37].

Here, we combined dense CRISPR-Cas9 mutagenesis with machine learning-guided functional analysis to systematically interrogate the WAVE regulatory complex in a microglia-relevant cellular model. This approach identified discrete functional regions within WRC subunits that either promote or suppress migration and revealed previously unrecognized therapeutic entry points within the complex. By integrating functional screening with computational drug prediction and experimental validation, we further identified clinically approved compounds capable of modulating WRC-dependent migration. These findings provide a domain-level map of WRC function and nominate actionable regulators of a microglial process implicated in Alzheimer’s disease.

## Results

### Myeloid cells express a paralog-selective WRC and support pooled functional interrogation of migration

To assess the suitability of THP-1 cells as a microglia-relevant myeloid model for investigating WRC function, we examined expression of WRC subunits using RNA-seq data generated from the same THP-1 cell population in our recently published study [24]. THP-1 cells robustly expressed the myeloid lineage marker *SPI1* (PU.1), confirming their myeloid identity. Most WRC components were expressed at moderate-to-high levels, including *BRK*1, *WASF2, CYFIP1, ABI1, ABI2*, and *NCKAP1L*. In contrast, several paralogous subunits were expressed at substantially lower levels, including *NCKAP1* relative to *NCKAP1L*, *CYFIP2* relative to *CYFIP1*, *WASF3* relative to *WASF1/WASF2*, and *ABI*3 relative to *ABI1/ABI2* (**Fig. 1A**) These data suggest that THP-1 cells predominantly assemble a WRC containing *NCKAP1L*, *CYFIP1*, and *WASF2*-family components, providing an experimentally tractable system for dissecting paralog-specific WRC function.

**Figure 1.**
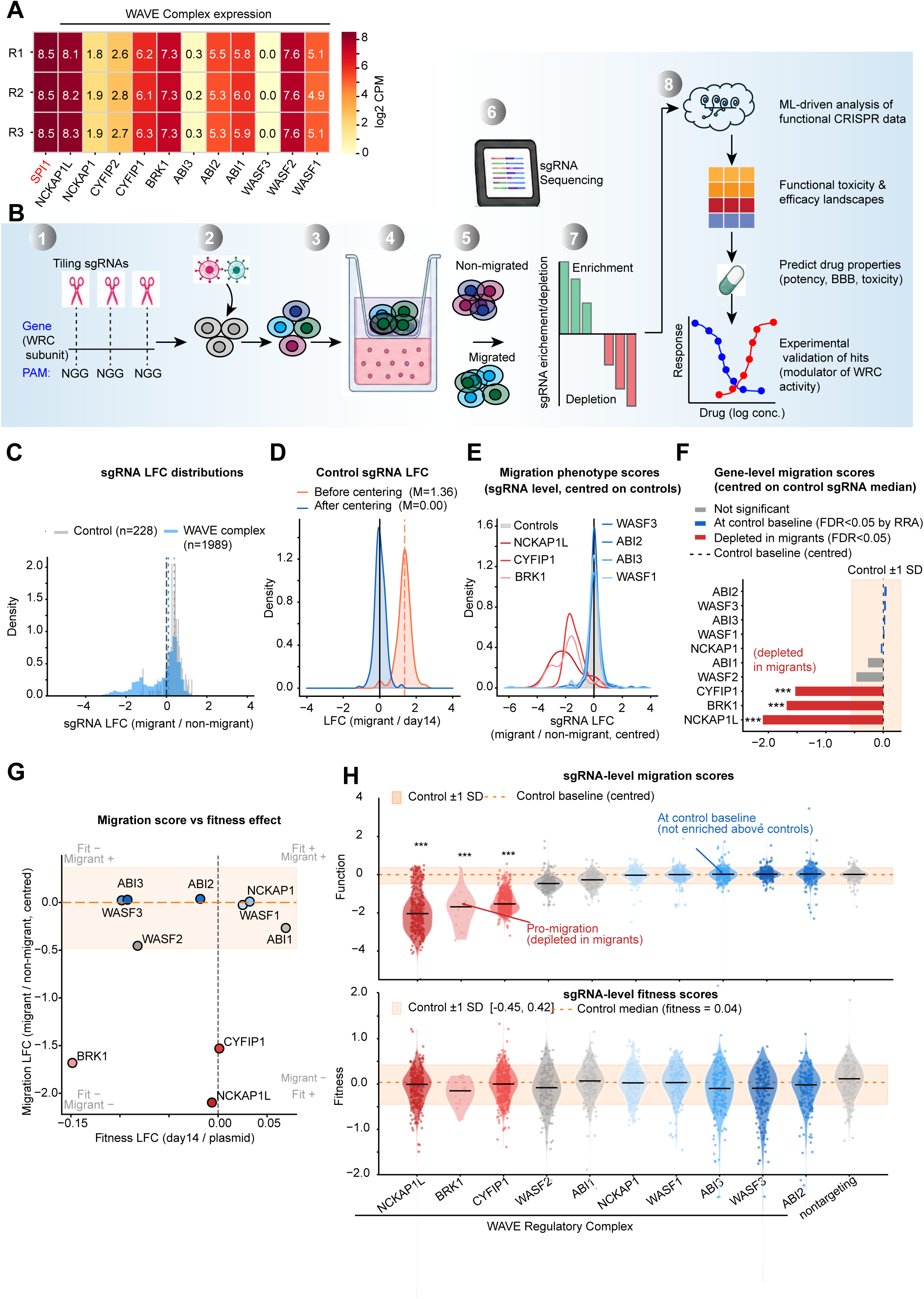
CRISPR tiling mutagenesis identifies WRC subunits that regulate CCL2-directed migration in THP-1 cells. **A**. Expression of WRC subunit genes and the myeloid marker *SPI1* in THP-1 cells, shown as log2 counts per million (CPM) from three biological replicates. **B**. Overview of the CRISPR tiling screen and CRISPRtile analysis workflow. A tiling sgRNA library targeting WRC subunits was introduced into Cas9-expressing THP-1 cells and subjected to a CCL2-directed transwell migration assay. sgRNA abundance in migrated and non-migrated populations was quantified by next-generation sequencing and analyzed using CRISPRtile to generate functional, fitness, and drug-prediction landscapes. **C**. Distribution of sgRNA-level migration scores (log2 fold change [LFC], migrant/non-migrant) for recovered WRC-targeting sgRNAs (n = 1,989 of 2,004 library sgRNAs) and control sgRNAs (n = 228). Fifteen WRC-targeting sgRNAs were not recovered following expansion and were excluded from downstream analyses. Dashed lines indicate medians. **D**. Distribution of control sgRNA migration LFCs before and after median-centering normalization. Control sgRNAs define the empirical baseline for normally migrating THP-1 cells in the CCL2 transwell assay, and median centering sets neutral migration behavior to zero for downstream analyses. **E**. Representative control-centered sgRNA-level migration score distributions for control sgRNAs and selected WRC subunits. sgRNAs targeting *NCKAP1L*, *CYFIP1*, and *BRK1* show depletion from the migrated fraction relative to the control baseline, whereas sgRNAs targeting other WRC paralogs remain near the control distribution. **F**. Gene-level control-centered migration scores, calculated as the median sgRNA migrated/non-migrated LFC for each gene. Red indicates significant depletion from the migrated population relative to controls; gray indicates no significant migration phenotype. ***, FDR < 0.05. **G**. Relationship between gene-level control-centered migration scores and fitness scores. Migration scores were calculated from migrated versus non-migrated fractions, whereas fitness scores were calculated from sgRNA representation after expansion relative to the plasmid library. **H**. sgRNA-level migration scores and fitness scores for each WRC subunit and control sgRNAs. *NCKAP1L*-, *BRK1*-, and *CYFIP1*-targeting sgRNAs are significantly depleted from the migrated fraction relative to control sgRNAs, identifying these subunits as required for efficient CCL2-directed migration. Statistical significance is shown relative to controls.

To systematically interrogate WRC-dependent migration, we designed a CRISPR-Cas9 tiling library spanning the coding sequences of 10 WRC subunit genes together with non-targeting and neutral-region controls (**Fig. S1A-B**). The final library contained 2,232 sgRNAs, including 2,004 guides targeting WRC genes (*ABI1, ABI2, ABI3, BRK1, CYFIP1, NCKAP1, NCKAP1L, WASF1, WASF2, WASF3*) and 228 control guides. The library was introduced into Cas9-expressing THP-1 cells and allowed to undergo seven population doublings prior to phenotypic selection. Cells were then subjected to a CCL2-directed transwell migration assay, and sgRNA abundance was quantified by next-generation sequencing in the plasmid library, the pre-migration population, and the migrated and non-migrated fractions (**Fig. 1B**). Because cells carrying non-targeting or neutral-region sgRNAs are expected to behave like unperturbed THP-1 cells, these control guides define the baseline distribution of normally migrating cells in the CCL2 transwell assay. In the absence of a migration phenotype, gene-targeting sgRNAs should distribute similarly to control sgRNAs between the migrated and non-migrated fractions. Initial analysis showed that the control-guide distribution was offset from zero, reflecting the empirical migrated/non-migrated baseline of normally migrating THP-1 cells in this assay (**Fig. 1D**). We therefore centered sgRNA migration scores on the median of the control-guide distribution, defining neutral migration behavior as zero for downstream analyses (**Fig. 1E)**

Together, these results established a robust screening framework for systematically identifying WRC components that promote or restrain CCL2-directed migration.

### NCKAP1L, BRK1, and CYFIP1 are required for efficient CCL2-directed migration

Gene-level analysis of control-centered migration scores identified three WRC subunits with significant depletion from the migrated population: *NCKAP1L*, *BRK1*, and *CYFIP1* (FDR < 0.05; **Fig. 1F**). These results indicate that perturbation of these subunits impairs efficient CCL2-directed migration and identify them as migration-required WRC components in THP-1 myeloid cells. In contrast, *ABI1*, *ABI2*, *ABI3*, *NCKAP1*, *WASF1*, *WASF2*, and *WASF3* did not show significant migration phenotypes after control centering. Notably, these non-significant genes were not enriched above the control baseline, indicating that their perturbation did not confer a reproducible migratory advantage relative to normally migrating control cells under the conditions tested. The three significant migration-required subunits corresponded to components expressed in THP-1 cells, with *NCKAP1L* representing the dominant *NCKAP* paralog and *CYFIP1* the dominant *CYFIP* paralog. This pattern suggests that the migration phenotype is concentrated in the WRC configuration most highly represented in this myeloid model rather than being uniformly distributed across all WRC paralogs.

To determine whether migration phenotypes reflected migration-specific effects or broader changes in cellular fitness, we compared control-centered migration scores with fitness scores derived from sgRNA abundance before and after cell expansion (**Fig. 1G**). *CYFIP1* displayed a strong migration phenotype with minimal fitness effect, indicating that *CYFIP1* perturbation selectively impaired migration rather than broadly compromising cell expansion. *NCKAP1L* and *BRK1* also showed strong depletion in the migrated fraction, with *BRK1* displaying a more evident negative fitness effect than *CYFIP1*. However, the magnitude and direction of the migration phenotypes were clearly distinguishable from the fitness baseline, supporting a primary role for these genes in migration.

Consistent with the gene-level analysis, sgRNA-level distributions confirmed that *NCKAP1L*-, *BRK1*-, and *CYFIP1*-targeting guides were significantly depleted from migrated cells relative to control sgRNAs (**Fig. 1H**). In contrast, sgRNAs targeting *ABI2*, *ABI3*, *NCKAP1*, *WASF1*, and *WASF3* remained near the control baseline and were not enriched above controls, further supporting the conclusion that *NCKAP1L*, *BRK1*, and *CYFIP1* are the most robust migration-required WRC subunits in this assay.

Together, these results identify *NCKAP1L*, *BRK1*, and *CYFIP1* as principal WRC components required for CCL2-directed migration in THP-1 myeloid cells. The separation between migration and fitness effects, particularly for *CYFIP1*, indicates that dense CRISPR mutagenesis can resolve WRC-dependent migration phenotypes that are not simply explained by impaired cell survival or proliferation. These findings nominate specific WRC subunits for residue-level functional mapping and pharmacological interrogation.

### CRISPRtile identifies discrete functional regions within NCKAP1L and CYFIP1

Among the migration-required WRC subunits identified in the primary screen, *NCKAP1L*, *CYFIP1*, and BRK1 showed the strongest phenotypes (**Fig. 1F**). Because *BRK1* is a small 75-amino-acid protein that affords limited resolution for Cas9 tiling analysis, we focused subsequent residue-level mapping on the two large core WRC subunits, *NCKAP1L* and *CYFIP1*. Dense tiling coverage was achieved for both genes (330 sgRNAs for *NCKAP1L* and 401 sgRNAs for *CYFIP1*; **Fig. S1B**), enabling systematic mapping of migration-associated functional landscapes. CRISPRtile-derived Function scores revealed highly non-uniform distributions of functional importance across both proteins (**Fig. 2A,B**). Rather than being evenly distributed, migration-associated functional residues clustered within discrete regions separated by extended segments with comparatively neutral scores. Similar patterns were observed after correction for guide-level biases, indicating that these features were not driven by local sgRNA effects. Additional per-residue annotations, including fitness (Dropout) scores, evolutionary conservation, and predicted structural features, are provided in **Fig. S2A,B**. Mapping Function scores onto AlphaFold [28] structural models demonstrated that the most critical residues clustered within defined structural regions (**Fig. 2C,D**). In *NCKAP1L*, high-magnitude Function scores localized primarily to two annotated domains spanning residues 12-185 and 807-1127. In *NCKAP1L*, high-magnitude Function scores localized primarily to two annotated domains spanning residues 12-185 and 807-1127. Notably, the N-terminal domain contains residues R129 and V141, where pathogenic *NCKAP1L* variants associated with IMD72 have been reported [9]. The p.R129W and p.V141F substitutions destabilize the WRC and impair immune-cell function, providing independent support for the biological relevance of the CRISPRtile-derived functional landscape. In CYFIP1, the dominant functional region localized to residues 702-910, corresponding to a high-confidence structural subdomain within the broader FragX_IP/Rac1-binding region.

**Figure 2.**
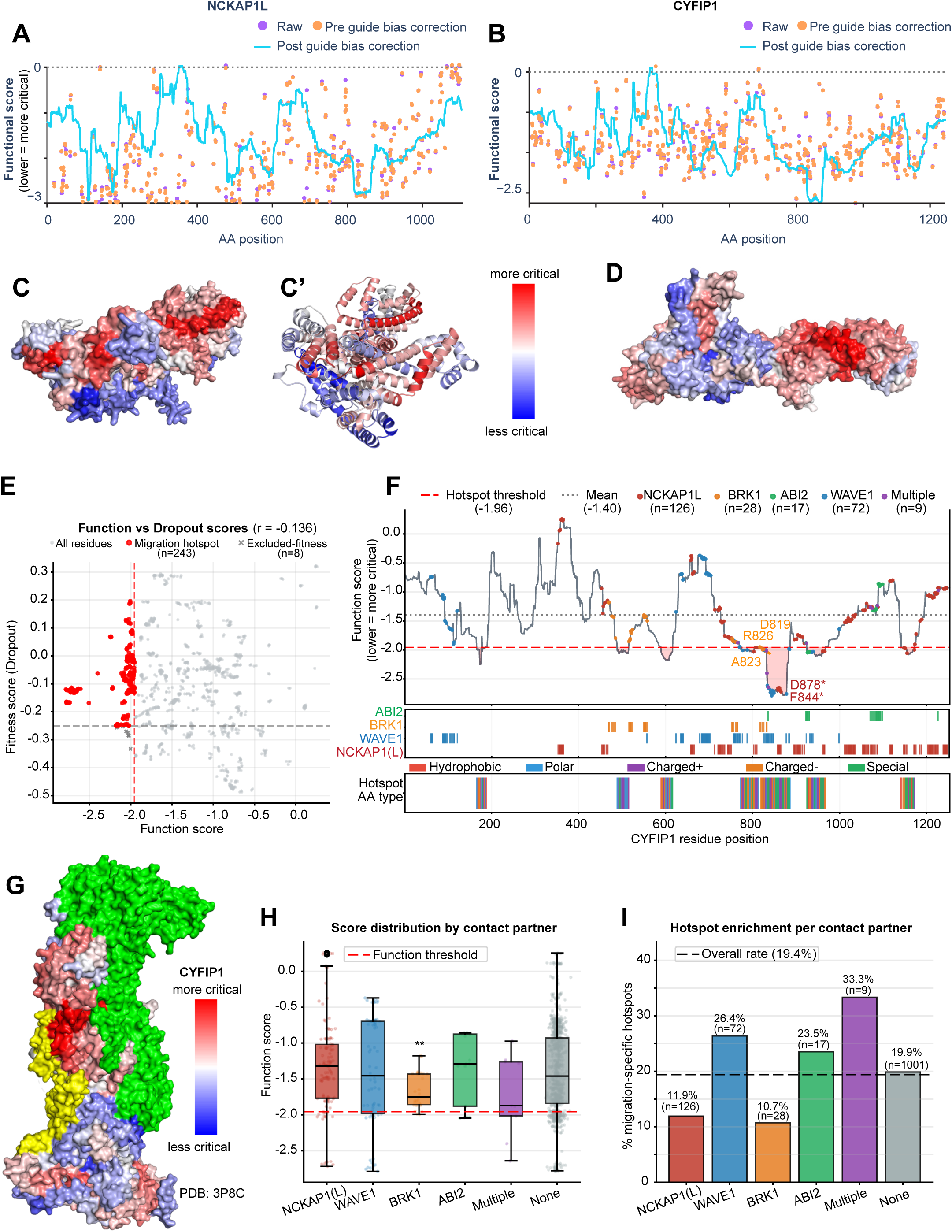
CRISPRtile identifies migration-associated functional regions and structural hotspots within NCKAP1L and CYFIP1. **A-B**. Per-residue CRISPRtile Function scores across the coding sequences of NCKAP1L (A) and CYFIP1 (B). Lower (more negative) scores indicate residues whose perturbation have a stronger impact on migration. Raw, guide-corrected, and final CRISPRtile prediction tracks are shown. **C-D**. Mapping of CRISPRtile Function scores onto AlphaFold structural models of NCKAP1L (C) and CYFIP1 (D). Residues are colored according to Function score, with more negative values indicating greater functional importance. **E**. Relationship between Function and Dropout (fitness) scores for all CYFIP1 residues. Red points indicate migration-specific residues (hotspot). Gray × symbols indicate residues excluded by the Dropout filter. Dashed lines indicate Function and Dropout thresholds used for hotspot classification. Pearson correlation coefficient (r) is shown. **F**. Identification of migration-specific hotspot residues in CYFIP1. Top: Function scores across the CYFIP1 sequence. Pink shaded regions indicate hotspot clusters defined by Function ≤ −1.96 and Dropout > −0.25. Colored circles denote residues making direct atomic contacts with NCKAP1(L), WAVE1, BRK1, ABI2, or multiple WRC partners in the 3P8C structure. Middle: Distribution of WRC contact residues along the CYFIP1 sequence. Bottom: Positions and physicochemical classes of migration-specific hotspot residues. **G**. Function scores mapped onto the crystal structure of the WRC pentamer (PDB: 3P8C). CYFIP1 is colored according to Function score. NCKAP1 (used as a structural proxy for NCKAP1L) is shown in green, BRK1 in yellow, and the remaining WRC subunits in gray. **H**. Distribution of Function scores for CYFIP1 residues grouped according to WRC contact partner. Box plots show median and interquartile range; points represent individual residues. Significance was assessed relative to residues lacking contacts with the indicated WRC partners using a one-sided Mann-Whitney U test. **, p < 0.01; ns, not significant. **I**. Frequency of migration-specific hotspot residues within each WRC contact category. The dashed line indicates the overall hotspot frequency across CYFIP1. Numbers above bars indicate the percentage of hotspot residues and total residue count for each category.

In contrast, fitness-critical residues were substantially more widespread throughout *NCKAP1L* than *CYFIP1* (**Fig. S2C,D**), consistent with the stronger fitness phenotype observed following *NCKAP1L* disruption in the pooled screen (**Fig. 1G,H**).

Together, these analyses demonstrate that migration-associated WRC function is concentrated within discrete structural regions rather than distributed uniformly across *NCKAP1L* and *CYFIP1*, nominating specific domains for downstream drug-target discovery and experimental validation.

### Migration-specific CYFIP1 hotspots localize to regulatory interfaces within the WRC

To identify residues specifically associated with migration rather than general cellular fitness, we combined CRISPRtile Function and Dropout scores. Residues were classified as migration-specific hotspots if they exhibited strongly negative Function scores (≤20th percentile; Function ≤ −1.96) while maintaining near-neutral fitness effects (Dropout > −0.25). Function and Dropout scores showed only weak correlation across all 1,253 CYFIP1 residues (Pearson r = −0.136; **Fig. 2E**), indicating that migration-associated functional importance is largely separable from effects on cellular fitness. Consistent with this observation, eight residues exhibited both strong migration and strong fitness phenotypes and were excluded from hotspot analysis. Application of these criteria identified 243 migration-specific hotspot residues, representing 19.4% of the CYFIP1 sequence.

To place migration-specific hotspots within the structural context of the WRC, we mapped CYFIP1 residues onto the crystal structure of the WRC pentamer (PDB : 3P8C) [6]. Application of the hotspot criteria identified multiple migration-associated regions distributed throughout the CYFIP1 sequence (**Fig. 2F and Fig. S2E**). However, these regions were not equivalent in magnitude. A single dominant hotspot spanning approximately residues 832-883 exhibited the most strongly negative CRISPRtile-Function scores across the protein, whereas the remaining hotspot regions clustered near the threshold used for hotspot classification. Mapping Function scores onto the assembled WRC revealed that functionally important residues were concentrated on a restricted surface of CYFIP1 rather than being dispersed throughout the structure (**Fig. 2G**). Notably, the dominant 832-883 hotspot overlapped the CYFIP1 interface with WAVE1 and was positioned adjacent to BRK1 and NCKAP1(L) contact surfaces. Residues F844 and D878, which make direct atomic contacts with WAVE1, delineated the boundaries of this hotspot, whereas D819, A823, and R826 localized near the neighboring BRK1 interaction surface. Together, these findings identify a prominent migration-associated functional region centered on the WAVE1/BRK1 interaction surface of CYFIP1.

We next asked whether residues contacting specific WRC subunits exhibited enhanced functional importance. Comparison of Function score distributions revealed that residues contacting BRK1 displayed significantly lower (more critical) Function scores than non-contact residues (one-sided Mann-Whitney U test, p = 0.008), whereas contacts involving NCKAP1L, WAVE1, ABI2, or multiple partners did not reach statistical significance (**Fig. 2H**). Analysis of hotspot frequencies showed that residues contacting multiple WRC partners (33.3%), WAVE1 (26.4%), and ABI2 (23.5%) were enriched above the overall hotspot rate of 19.4%, whereas NCKAP1L-contacting residues were depleted (11.9%) (**Fig. 2I**). Although BRK1-contact residues exhibited a below-average hotspot frequency (10.7%), their overall distribution was shifted toward more negative Function scores, indicating a broad contribution of the BRK1 interaction surface to migration-associated CYFIP1 activity rather than the presence of a small number of highly enriched hotspot residues.

Collectively, these analyses identify discrete migration-specific functional regions within CYFIP1 and localize them to regulatory interfaces within the WRC. The dominant hotspot cluster centered on residues 820-880 emerged as a particularly attractive candidate for therapeutic targeting because of its strong migration phenotype, minimal fitness effects, and strategic location at the interface between CYFIP1 and neighboring WRC subunits.

### CRISPRtile-guided drug prediction identifies Montelukast sodium as a modulator of CYFIP1-dependent migration

Having identified a dominant migration-associated hotspot within CYFIP1 (residues 832-883; **Fig. 2**), we next asked whether this region could be exploited for function-based drug discovery. To identify compounds predicted to interact with this hotspot, we screened a library of 3,229 compounds using TransformerCPI2.0 [5], a transformer-based compound-protein interaction model integrated within the CRISPRtile pipeline. The target region encompassed the major hotspot cluster identified by CRISPR tiling together with adjacent BRK1-associated residues implicated in migration regulation (Fig. 3A-D).

**Figure 3.**
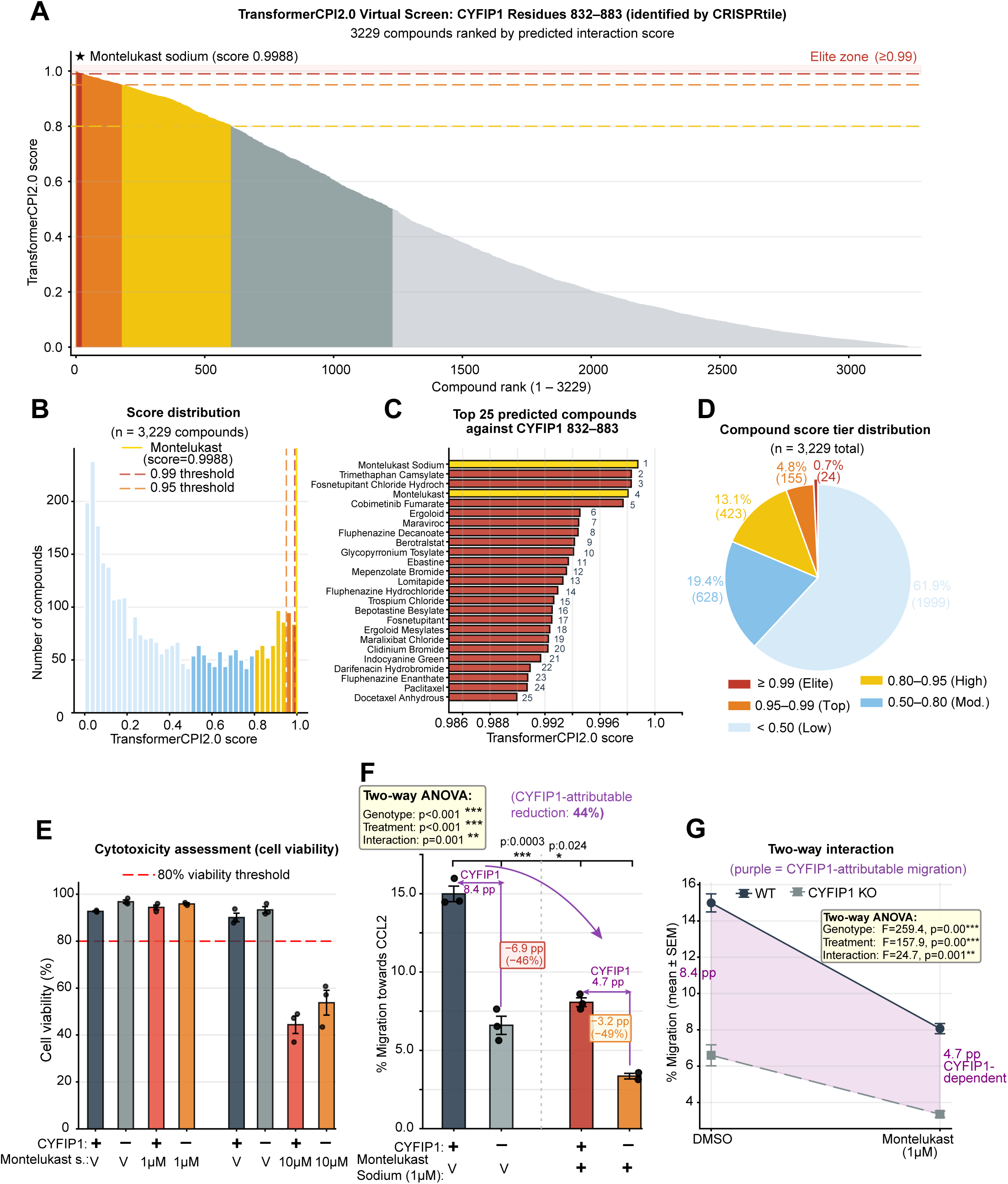
CRISPRtile-guided drug prediction identifies Montelukast sodium as a modulator of CYFIP1-dependent migration. **A**. Ranked distribution of TransformerCPI2.0 interaction scores for 3,229 compounds screened against the CYFIP1 hotspot region (residues 832-883). Compounds are ordered by rank, with colors indicating score tiers. Montelukast sodium (star) ranked first with a score of 0.9988. **B.** Distribution of TransformerCPI2.0 interaction scores across the screened compound library. Vertical lines indicate elite-tier (≥0.99) and top-tier (≥0.95) thresholds. Montelukast sodium is indicated by the gold line. **C**. Top 25 compounds ranked by TransformerCPI2.0 interaction score. Montelukast sodium is highlighted in gold. **D**. Proportion of screened compounds within each TransformerCPI2.0 score tier. The elite tier (≥0.99) comprised 24 compounds (0.74% of the screened library). **E.** Cell viability of WT and *CYFIP1* KO THP-1 cells following treatment with Montelukast sodium (1 µM and 10 µM). Values represent mean ± SEM (n = 3). **F**. CCL2-directed transwell migration of WT and *CYFIP1* KO THP-1 cells following treatment with vehicle (DMSO) or Montelukast sodium (1 µM). Bars represent mean ± SEM with individual biological replicates overlaid. Purple arrows indicate the CYFIP1-attributable migration component (WT − *CYFIP1* KO) under each treatment condition. Statistical comparisons between vehicle- and drug-treated groups were performed using two-sided t-tests. Two-way ANOVA results are shown in the inset. One *CYFIP1* KO + Montelukast replicate was excluded because viability fell below the predefined assay threshold. **G**. Two-way ANOVA interaction plot showing migration responses of WT and *CYFIP1* KO THP-1 cells following vehicle or Montelukast treatment. Shaded regions indicate the CYFIP1-attributable migration component under each condition. Insets show ANOVA statistics for genotype, treatment, and genotype × treatment interaction effects.

TransformerCPI2.0 assigned interaction scores ranging from 0 to 1, with higher scores indicating stronger predicted interactions with the CYFIP1 hotspot. The resulting score distribution was strongly bimodal, indicating clear separation between predicted interactors and non-interactors (Fig. 3A-D). Only 24 compounds (0.74% of the screened library) achieved elite-tier scores (≥0.99). Among all compounds tested, Montelukast sodium, an FDA-approved leukotriene receptor antagonist with reported blood-brain barrier penetration [44, 56], ranked first with a predicted interaction score of 0.9988 (Fig. 3A-D).

To determine whether Montelukast sodium modulates WRC-dependent migration, we performed CCL2-directed transwell migration assays using wild-type (WT) and *CYFIP1* knockout (KO) THP-1 cells. Cells were pre-treated with vehicle (DMSO) or Montelukast sodium prior to migration and quantified using multiple independent migration metrics (**Fig. 3F,G; Fig. S3A-F**). At 1 µM, Montelukast sodium maintained high cell viability across all conditions (>93%; Fig. 3E), indicating that migration phenotypes were not attributable to overt cytotoxicity.

Montelukast sodium significantly reduced migration of WT THP-1 cells from 15.0% to 8.1% (−46.2%; p = 0.0003) (Fig. 3F). Migration of *CYFIP1* KO cells was also reduced, from 6.6% to 3.4% (−49.1%; p = 0.024), although the absolute magnitude of inhibition was substantially smaller than that observed in WT cells. Consistent with this observation, two-way ANOVA identified significant effects of genotype (p < 0.0001), treatment (p < 0.0001), and genotype × treatment interaction (p = 0.001), indicating that the response to Montelukast sodium was partially dependent on *CYFIP1* status (Fig. 3F-G).

To estimate the CYFIP1-dependent component of migration, we compared the difference in migration between WT and *CYFIP1* KO cells under each treatment condition. Montelukast sodium reduced this CYFIP1-attributable migratory component from 8.4 to 4.7 percentage points (PP), representing an approximately 44% reduction in CYFIP1-dependent migration (Fig. 3F,G). Residual inhibition observed in *CYFIP1* KO cells is consistent with additional CYFIP1-independent effects arising from the established pharmacology of Montelukast sodium. At 10 µM, Montelukast sodium substantially reduced cell viability and therefore precluded meaningful interpretation of migration phenotypes (Fig. 3E). Consequently, all mechanistic conclusions were based on the 1 µM condition.

Collectively, these findings demonstrate that functionally important regions identified by CRISPR tiling can be exploited pharmacologically to modulate WRC-dependent migration, validating CYFIP1 as a proof-of-concept target for hotspot-guided therapeutic discovery.

### NCKAP1L is a microglia-enriched WRC component broadly expressed across microglial states

To determine the relevance of migration-regulating WRC subunits in the human brain, we examined their expression in published single-nucleus RNA-sequencing datasets [20, 54] (Fig. 4A-B and Fig. S4). Among the migration-promoting subunits identified in our screen, *CYFIP1* was broadly expressed across multiple brain cell populations, including excitatory neurons, inhibitory neurons, astrocytes, vascular cells, and microglia. In contrast, *NCKAP1L* expression was highly enriched in microglia, with minimal expression detected in other major brain cell types (Fig. 4A-B). *BRK1* showed detectable expression in microglia but was also expressed in additional cell populations, whereas *WASF1* was predominantly neuronal. These data identify *NCKAP1L* as the most microglia-restricted migration-promoting WRC component emerging from our functional screen.

**Figure 4.**
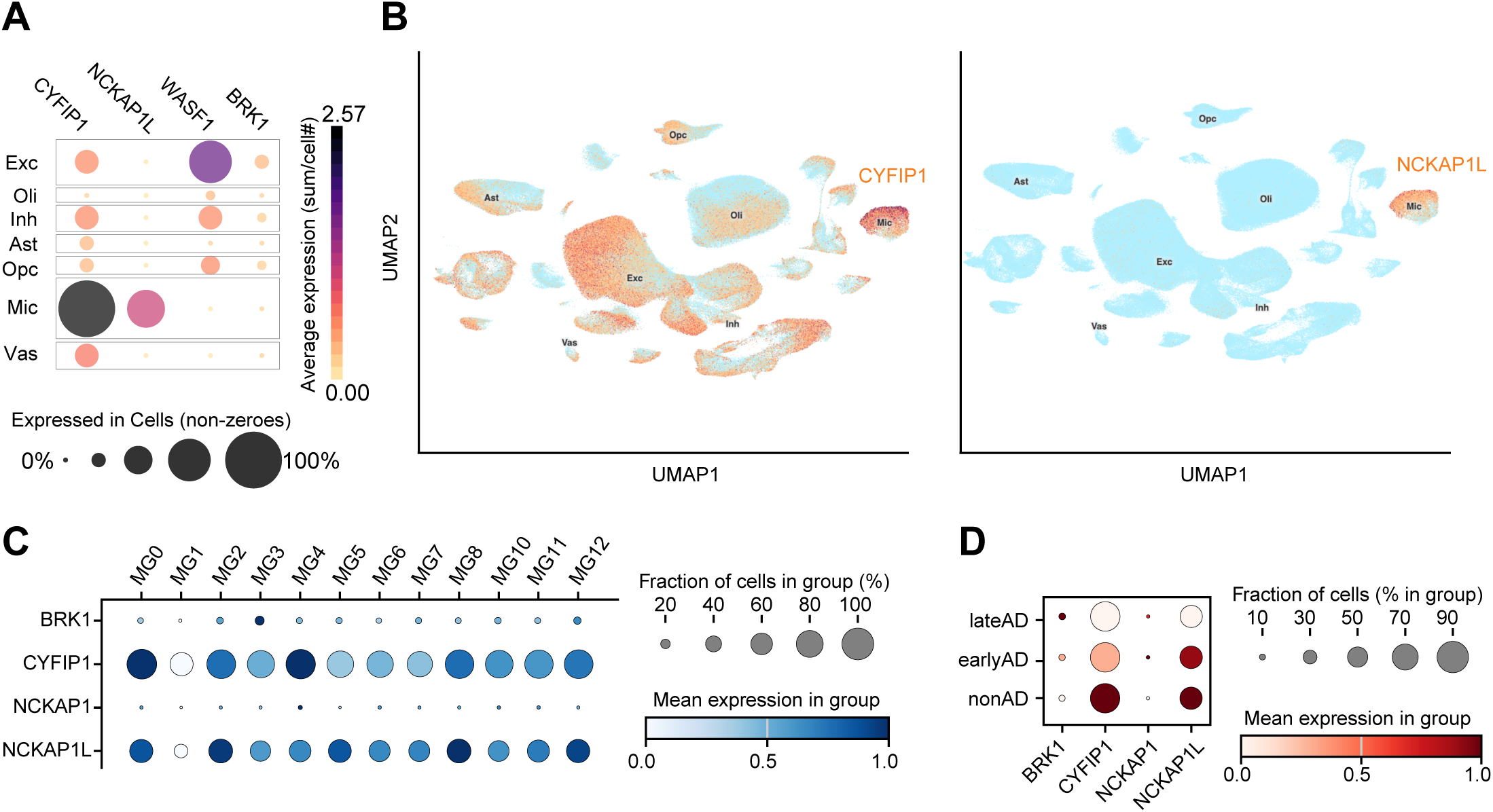
NCKAP1L is a microglia-enriched WRC component broadly expressed across microglial states. **A**. Expression of migration-associated WRC subunits across major human brain cell types in public single-nucleus RNA-sequencing data [54] from human prefrontal cortex. Dot size indicates the fraction of cells expressing each gene, and color indicates average expression level. Exc, excitatory neurons; Inh, inhibitory neurons; Ast, astrocytes; Oli, oligodendrocytes; Opc, oligodendrocyte precursor cells; Mic, microglia; Vas, vascular cells. **B**. UMAP visualization of *CYFIP1* and *NCKAP1L* expression across human brain cell populations. Cells are colored according to normalized gene expression. *NCKAP1L* expression is highly restricted to microglia, whereas *CYFIP1* is broadly expressed across multiple cell types. **C**. Expression of *BRK1, CYFIP1, NCKAP1*, and *NCKAP1L* across twelve transcriptionally defined human microglial states. Dot size represents the fraction of cells expressing each gene and color indicates mean expression level within each microglial state. **D**. Expression of *BRK1, CYFIP1, NCKAP1*, and *NCKAP1L* in microglia from non-AD, early-AD, and late-AD individuals. Dot size indicates the fraction of expressing cells and color represents mean expression within each disease stage.

To further characterize *NCKAP1L* expression within the microglial compartment, we analyzed single-nucleus transcriptomic data [51] spanning twelve human microglial states associated with homeostatic (MG0), inflammatory (MG2, MG8, MG10), phagocytic (MG5), lipid-processing (MG4), glycolytic (MG7), antiviral (MG11), and proliferative programs (MG12) (Fig. 4C). *NCKAP1L* was robustly expressed across most of microglial states, whereas *NCKAP1* expression remained uniformly low. *CYFIP1* was likewise broadly expressed across microglial populations, while *BRK1* expression was comparatively modest. The widespread expression of *NCKAP1L* across diverse microglial states suggests that it represents a core component of the microglial transcriptional program rather than a marker restricted to a specific activation state.

We next examined expression of migration-promoting WRC subunits across microglia from non-AD, early-AD, and late-AD individuals (Fig. 4D). While expression levels varied modestly across disease stages, differential expression analysis did not identify consistent disease-associated changes for *CYFIP1, NCKAP1,* or *NCKAP1L*. These analyses suggest that NCKAP1L expression is maintained throughout AD progression and reinforce its role as a core component of the microglial transcriptional program rather than a disease-stage-specific marker. Given its microglia-restricted expression, robust expression across diverse microglial states, and strong requirement for chemokine-directed migration, we further prioritized NCKAP1L for subsequent functional mapping and therapeutic interrogation.

### NCKAP1L contains a conserved migration-associated hotspot with features compatible with pharmacological targeting

Having established *NCKAP1L* as a microglia-enriched regulator of migration (**Figs. 1 and 4**), we next sought to identify discrete NCKAP1L regions suitable for therapeutic modulation. Consistent with a biologically meaningful functional landscape, CRISPRtile Function scores showed a significant relationship with evolutionary conservation, with residues exhibiting stronger migration phenotypes tending to be more highly conserved (Pearson r = 0.30, p = 3.38 × 10^-28^; **Fig. 5A**). Function and fitness scores remained largely separable (**Fig. 5B**), indicating that many residues contributing to migration could be distinguished from those broadly affecting cellular fitness. To further prioritize candidate target sites, we integrated CRISPRtile Function scores with evolutionary conservation, secondary structure annotation, and solvent accessibility (**Fig. 5C**). This analysis identified several residues exhibiting strongly negative Function scores together with structural accessibility, suggesting the presence of pharmacologically tractable surfaces within NCKAP1L. To define candidate functional segments in an unbiased manner, we applied the TGUH algorithm from ProTiler [25], a computational framework originally developed to identify CRISPR knockout hypersensitive (CKHS) protein regions from tiling-sgRNA knockout screens . Here, we applied this region-calling strategy to guide-efficiency-corrected CRISPRtile Function scores, thereby identifying NCKAP1L segments in which CRISPR-induced perturbations produced strong migration phenotypes. This analysis identified five functionally hypersensitive NCKAP1L regions spanning residues 91-121, 133-194, 410-429, 483-624, and 719-1027 (**Fig. 5D**).

**Figure 5.**
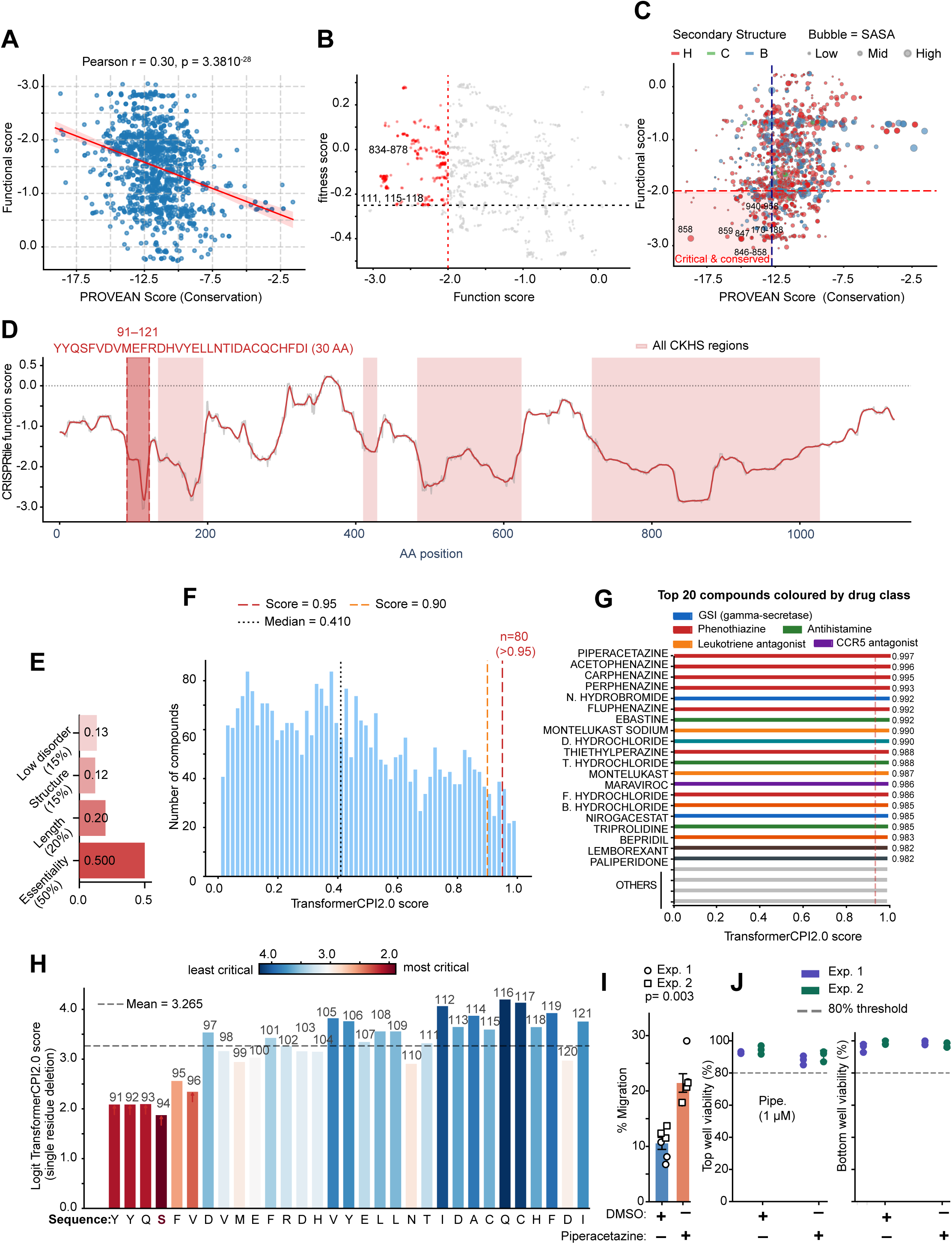
CRISPRtile prioritizes an NCKAP1L regulatory region and identifies Piperacetazine as a positive modulator of migration. **A**. Relationship between CRISPRtile Function scores and evolutionary conservation across NCKAP1L residues. Each point represents one amino acid residue. More negative Function scores indicate stronger contribution to CCL2-directed migration, whereas more negative PROVEAN scores indicate greater evolutionary constraint. Pearson correlation coefficient and corresponding p value are shown. **B**. Relationship between CRISPRtile Function scores and Dropout (fitness) scores across NCKAP1L residues. Each point represents one amino acid residue. Red points indicate candidate migration-associated residues with strong Function effects and limited Dropout effects, representing potential migration-specific residues for downstream prioritization. **C**. Integrated assessment of residue criticality, conservation, and solvent accessibility in NCKAP1L. Each bubble represents one residue plotted by PROVEAN score and CRISPRtile Function score. Bubble size indicates solvent-accessible surface area (SASA), and color indicates secondary structure assignment: helix (H), coil (C), or bridge (B). Dashed lines indicate the bottom 25th percentile thresholds for Function and PROVEAN scores; the shaded region highlights residues that are both functionally critical and evolutionarily conserved. Selected residues are annotated. **D**. CRISPRtile Function score profile and CRISPR knockout hypersensitive-like region annotation across NCKAP1L. The gray line shows guide-efficiency-corrected Function scores across the full protein, and the colored line shows a 7-residue smoothed profile. Shaded regions indicate functionally hypersensitive segments identified using TGUH segmentation. The selected region used for compound-interaction prediction, residues 91-121, is highlighted. **E**. Composite prioritization of NCKAP1L candidate regions for drug screening. Horizontal bars show weighted contributions of functional importance, target length, structural order, and low intrinsic disorder to the final composite score for the selected NCKAP1L region. **F**. TransformerCPI2.0 virtual screen of 3,229 compounds against the prioritized NCKAP1L 91-121 region. The distribution of predicted interaction scores is shown, with dashed lines indicating 0.90 and 0.95 score thresholds. Eighty compounds (2.5%) exceeded the 0.95 threshold. **G**. Top-scoring compounds from the NCKAP1L 91-121 virtual screen. Bars show TransformerCPI2.0 interaction scores for the top 20 compounds, colored by pharmacological class. Piperacetazine was the highest-scoring compound. Compound structures were represented using SMILES strings retrieved from ChEMBL. **H**. In silico single-residue deletion mutagenesis of the NCKAP1L 91-121 region using Piperacetazine as the query compound. Each bar represents the predicted interaction score after deletion of the indicated residue. Lower scores indicate greater predicted contribution to compound interaction. Residues are colored according to relative importance, with red indicating the strongest predicted contribution. S94 produced the largest reduction in predicted interaction score. **I.** CCL2-directed transwell migration of THP-1 cells treated with vehicle (DMSO) or Piperacetazine (1 µM). Migration is shown as percentage of cells recovered from the lower chamber. Data represent six replicates per condition across two independent experiments. **J**. Viability of THP-1 cells from the transwell migration assay shown in panel I. Viability was assessed in upper and lower chamber populations. The dotted line indicates the 80% assay-validity threshold.

Because multiple functionally hypersensitive NCKAP1L regions were identified, we next applied a composite prioritization framework to select the region most suitable for downstream drug screening. This framework integrated CRISPR-derived functional importance with features relevant to pharmacological tractability, including target length, structural order, and low intrinsic disorder. Among the five candidate regions, residues 91-121 achieved the highest composite score (0.954; Fig. 5E). This 30-amino-acid segment combined the strongest migration-associated functional signal with favorable structural properties, including high predicted order and minimal disorder. Notably, analogous analyses of CYFIP1 and BRK1 identified candidate regions that overlapped migration-associated hotspots detected in earlier analyses (**Fig. S5A-D**), supporting the robustness of the prioritization strategy. Together, these results nominate NCKAP1L residues 91-121 as a candidate regulatory surface for pharmacological modulation of WRC-dependent migration.

### Machine-learning-guided screening identifies Piperacetazine as a positive modulator of migration

To identify compounds predicted to modulate the prioritized NCKAP1L region, we screened 3,229 FDA-approved compounds against the NCKAP1L 91-121 sequence using the TransformerCPI2.0 module within CRISPRtile. Compound interaction scores displayed a bimodal distribution, with most compounds scoring below 0.7 and a restricted high-scoring tail (**Fig. 5F**). Eighty compounds (2.5% of the screened library) achieved scores greater than 0.95, indicating selective rather than broadly promiscuous predicted interactions.

Phenothiazine-class compounds were strongly represented among the highest-ranking candidates (**Fig. 5G**; **Fig. S5E**). The top-scoring compound was Piperacetazine (TransformerCPI2.0 score = 0.9972), followed by additional phenothiazines and several mechanistically distinct compounds, including Nirogacestat, Ebastine, Montelukast, and Maraviroc. The recovery of compounds previously linked to immune-cell migration or inflammatory signaling provided supportive evidence that the screen captured biologically relevant pharmacology.

To investigate the predicted interaction surface in greater detail, we performed in silico single-residue deletion mutagenesis across the NCKAP1L 91-121 region using Piperacetazine as the query compound. Deletion of residues within the N-terminal portion of the sequence produced the largest reductions in predicted interaction score, identifying the YYQSFV sequence spanning residues 91-96 as the principal predicted determinant of compound recognition (**Fig. 5H**). Among these residues, S94 produced the strongest reduction in predicted interaction score upon deletion, suggesting that the N-terminal face of this region contributes disproportionately to the predicted Piperacetazine interaction.

Finally, we experimentally tested Piperacetazine in CCL2-directed transwell migration assays. Treatment of THP-1 cells with 1 µM Piperacetazine significantly increased migration relative to vehicle-treated (DMSO) controls, with all replicates showing a consistent directional response (**Fig. 5I**; **Fig. S5G,H**). Cell viability remained above assay-quality thresholds and was not significantly affected by treatment (**Fig. 5J**), indicating that the increased migration was not attributable to altered cell survival. Together, these findings identify Piperacetazine as a positive modulator of migration and support the NCKAP1L 91-121 region, positioned at the NCKAP1L-CYFIP1 interface within the WRC structure, as a candidate regulatory surface for pharmacological modulation of WRC-dependent migration.

## Discussion

Microglia have emerged as central cellular mediators of Alzheimer’s disease (AD) risk, progression, and tissue response to pathology. Genetic and epigenomic studies have repeatedly shown that many AD-associated risk loci map to regulatory elements or genes enriched in microglia, supporting the view that microglial biology contributes to disease susceptibility rather than simply reflecting a downstream reaction to amyloid-β and tau pathology [24, 30, 41, 54]. However, many AD risk genes remain difficult to translate into therapeutic mechanisms because their disease-relevant cellular functions and druggable protein regions are unknown. In this study, we addressed this problem by applying dense CRISPR-Cas9 mutagenesis and CRISPRtile-based functional mapping to the WAVE regulatory complex (WRC), an actin-remodeling complex positioned at the interface between extracellular cues, cytoskeletal remodeling, and immune-cell migration.

The WRC is particularly relevant to AD because microglial migration and lesion-directed positioning are essential components of the brain’s response to neurodegenerative pathology. Microglia must sense, migrate toward, and organize around amyloid plaques and injured tissue, where they can influence plaque compaction, phagocytosis, inflammatory signaling, synaptic remodeling, and tissue repair [2, 11, 59]. Consistent with this biology, deletion of the AD-risk gene ABI3, a WRC-associated component, exacerbates amyloid pathology and reduces microglial clustering around plaques in a 5XFAD model, linking WRC-associated actin regulation to microglial behavior in AD-relevant pathology [29]. Our data extend this concept by showing that WRC-dependent regulation of migration is not confined to *ABI3*-associated biology, but includes additional WRC components, particularly *NCKAP1L*, *CYFIP1*, and *BRK1*, that are required for efficient chemokine-directed myeloid migration and can be resolved further at the level of specific functional regions.

A key finding from near saturation CRISPR-screen analysis was that WRC subunits do not contribute equally to chemokine-directed migration. Loss of *NCKAP1L*, *CYFIP1*, and *BRK1* significantly reduced CCL2-directed migration, identifying these subunits as the principal migration-required WRC components in myeloid cells. In contrast, perturbation of *ABI1*, *ABI2*, *ABI3*, *NCKAP1*, *WASF1*, *WASF2*, and *WASF3* did not produce reproducible migration phenotypes relative to normally migrating control cells after control centering. Thus, the migration requirement was concentrated in a subset of WRC components rather than distributed uniformly across all paralogs. This pattern is consistent with the paralog-selective WRC expression program observed in THP-1 cells and suggests that the dominant WRC configuration in this myeloid model is particularly important for migration. Importantly, the CYFIP1 migration phenotype was largely separable from general fitness effects, whereas *NCKAP1L* and *BRK1* showed stronger accompanying fitness effects. This distinction is essential for therapeutic interpretation, because a useful modulator of myeloid or microglial migration should ideally alter motility without broadly compromising cell viability.

Another key aspect of this work is the demonstration that dense CRISPR mutagenesis can be integrated with machine-learning-based functional mapping to identify candidate pharmacological regulatory surfaces within large protein complexes. Conventional CRISPR knockout screens are powerful for identifying genes required for a phenotype, but they generally do not reveal which residues, domains, or protein surfaces mediate that phenotype. This limitation is particularly important for multiprotein complexes such as the WRC, where different subunits and interfaces can have distinct effects on complex stability, regulatory state, and cellular behavior. Building on our prior development of CRISPR tiling and computational frameworks for high-resolution functional mapping [3, 26, 43, 46]. CRISPRtile [37] extends this strategy to the AD-relevant problem of microglial migration by converting dense WRC mutagenesis data into residue-level functional and fitness landscapes. The key conceptual advance is that these landscapes can nominate actionable protein regions within a microglial pathway, creating a direct bridge from CRISPR perturbation data to therapeutic hypothesis generation.

Among the WRC subunits identified in the screen, *CYFIP1* provided a proof-of-principle example of hotspot-guided pharmacological modulation. CRISPRtile identified a dominant CYFIP1 migration-associated hotspot centered near residues 832-883, positioned at a WRC interface involving WAVE1 and BRK1. Drug prediction nominated Montelukast sodium, an approved cysteinyl leukotriene receptor antagonist, as a top-scoring compound for this region. Montelukast significantly reduced CCL2-directed migration, and the magnitude of inhibition was attenuated in *CYFIP1* knockout cells, supporting partial dependence on CYFIP1-associated pathways. This finding is notable because Montelukast has already attracted interest in neurodegeneration and AD, where leukotriene signaling has been linked to neuroinflammation, microglial activation, blood-brain barrier function, and cognition [22, 34, 44]. Our results do not prove direct binding of Montelukast to CYFIP1, but they raise the possibility that some effects of leukotriene-targeted pharmacology on immune-cell behavior may intersect with WRC-dependent cytoskeletal regulation.

*NCKAP1L* emerged as the most AD-relevant WRC subunit for follow-up because it combines a strong migration phenotype with marked microglial enrichment in human brain datasets. Unlike *CYFIP1*, which is broadly expressed across neural and glial cell types, *NCKAP1L* was highly enriched in microglia and expressed across diverse microglial transcriptional states. This pattern suggests that NCKAP1L is not simply a disease-stage marker, but rather a core component of the human microglial cytoskeletal program. This interpretation is consistent with human immunology studies showing that NCKAP1L, also known as HEM1, is a hematopoietic WRC component required for immune-cell function; pathogenic *NCKAP1L* variants, including p.R129W and p.V141F, destabilize the WRC and cause immunodeficiency 72 with autoinflammation and lymphoproliferation [4, 9]. The overlap between NCKAP1L disease-associated regions and CRISPRtile-defined functional regions supports the biological relevance of the migration landscape inferred in our screen.

The NCKAP1L analysis further demonstrates how functional maps can be converted into therapeutic hypotheses. Rather than selecting the most essential region alone, we prioritized candidate regions using a composite strategy that incorporated CRISPR-derived functional importance, target length, structural order, and low intrinsic disorder. This approach identified residues 91-121 as a candidate pharmacological regulatory region. Screening approved compounds against this sequence nominated Piperacetazine as the top-scoring compound, and computational mutagenesis localized predicted binding sensitivity to the N-terminal portion of this segment. Experimentally, Piperacetazine increased CCL2-directed migration without reducing viability, identifying it as a positive modulator of myeloid-cell migration. Together with the Montelukast result, these findings suggest that WRC-dependent migration can be pharmacologically tuned in both directions, providing a potential strategy for either dampening excessive inflammatory migration or enhancing lesion-directed microglial responses depending on disease context. This bidirectional control may be particularly relevant to AD, where microglial activity is unlikely to be uniformly beneficial or harmful. In early disease stages, efficient microglial surveillance, migration, and plaque engagement may help contain pathology, whereas chronic or mislocalized inflammatory responses may contribute to neurotoxicity at later stages [19, 53, 59]. Therefore, the therapeutic goal may not be global microglial suppression, but context-dependent tuning of specific microglial functions. Our data nominate WRC-mediated migration as one such tunable function. In this framework, CYFIP1- and NCKAP1L-associated regulatory regions represent potential entry points for adjusting microglial motility without broadly ablating microglial identity or viability.

Several limitations should be considered. First, the primary functional screen was performed in THP-1 cells, a tractable human myeloid model that does not fully recapitulate primary or iPSC-derived human microglia. Validation in human iPSC-derived microglia and, ultimately, disease-relevant AD model systems will be necessary to establish translational relevance. Second, the drug predictions and computational mutagenesis nominate candidate compound-protein interactions, but direct biochemical binding has not yet been demonstrated. Future studies using targeted mutagenesis, cellular thermal shift assays, biophysical binding assays, and structural approaches will be needed to define the molecular mechanisms of Montelukast and Piperacetazine action. Third, migration is only one microglial function. Whether WRC modulation also affects phagocytosis, inflammatory signaling, synaptic pruning, plaque engagement, or disease-associated microglial state transitions remains to be determined.

In summary, this study provides a sequence-resolved functional map of the WRC in a microglia-relevant migration assay and identifies CYFIP1 and NCKAP1L as pharmacologically actionable regulators of chemokine-directed migration. By integrating dense CRISPR mutagenesis, ML-based functional mapping, structural prioritization, and experimental drug validation, we establish a framework for converting AD-relevant microglial pathways into testable therapeutic hypotheses. These findings support WRC-dependent cytoskeletal regulation as an underexplored mechanism in AD biology and nominate microglial migration as a functionally tunable process for future therapeutic investigation.

## Acknowledgements

We thank the members of the Center for Translational & Computational Neuroimmunology and the Taub Institute for Research on Alzheimer’s Disease and the Aging Brain for technical assistance and helpful discussions.

## Data Availability

All processed data required to reproduce the main findings are provided in the Supplementary Tables. Supplementary Table 1 contains the WRC sgRNA library annotations used in this study.

Supplementary Table 2 contains sgRNA count tables, migration and fitness log2 fold-change values, and input files used for CRISPRtile analysis of the WRC migration assay. Supplementary Table 3 contains CRISPRtile output files, residue-level Function and Dropout scores, hotspot annotations, compound-prediction results, and master analysis files derived from the WRC screening data.

Raw sgRNA amplicon sequencing FASTQ files will be deposited in NCBI GEO/SRA prior to journal submission. All processed data required to reproduce the figures in this preprint are included in Supplementary Tables 1-3.

Public human single-nucleus and single-cell RNA-seq datasets analyzed in this study were obtained from previously published sources as described in Methods.

## Code Availability

No new code was generated in this study. The CRISPRtile [37] platform is available as a Google Colab at: https://colab.research.google.com/drive/14fvBFYtHNU73_JlbyRMZYwv8L84HKisk?usp=sharing All source code used in CRISPRtile are available from GitHub at :https://github.com/jasoncngo/CRISPRtile

## Author Contributions

Conceptualization, Methodology, and Investigation: J.C.N. and F.S.; Data Curation and Formal Analysis: J.C.N., Y.X., M.O., and F.S.; Writing, Review & Editing: F.S.; Funding Acquisition and Resources: F.S.; Supervision: F.S.

## Methods

### Cell culture

Human THP-1 monocytic cells were obtained from ATCC (TIB-202) and cultured in RPMI 1640 medium supplemented with 10% fetal bovine serum and 1% penicillin–streptomycin according to the supplier’s recommendations. HEK293T cells used for lentivirus production were cultured in Dulbecco’s modified Eagle’s medium supplemented with 10% fetal bovine serum and penicillin-streptomycin. Cells were maintained at 37°C in a humidified incubator with 5% CO_2_.

### Design and cloning of tiled sgRNA library

Tiled sgRNA libraries were designed by identifying all 20 nt sequences upstream of an NGG PAM on both sense and antisense strands across coding regions of WAVE regulatory complex genes. The library targeted WASF1, WASF2, WASF3, ABI1, ABI2, ABI3, BRK1, CYFIP1, NCKAP1, and NCKAP1L, together with non-targeting and genomic control sgRNAs. The final library contained 2,232 sgRNAs, including 2,004 WRC-targeting sgRNAs across 10 genes and 228 control sgRNAs. Oligonucleotide pools were synthesized on CustomArray Oligo (GenScript Biotech) and amplified by PCR. A second PCR was performed to remove synthesis barcodes and append vector-homology sequences. The amplified sgRNA library was cloned into LentiGuide-Puro (Addgene plasmid 52963) [42] using Gibson assembly. Library transformation was performed at sufficient scale to maintain approximately 1,800 bacterial colonies per sgRNA. The plasmid library was deep sequenced to confirm sgRNA representation.

### Lentivirus production

Lentivirus was produced in HEK293T cells cultured in 15 cm tissue-culture dishes. Cells were transfected at 70-80% confluence with 20 µg lentiviral sgRNA plasmid, 13.3 µg psPAX2 packaging plasmid (Addgene plasmid 12260), and 6.7 µg VSV-G envelope plasmid (Addgene plasmid 8454) using 180 µg linear polyethylenimine. Medium was changed 24 h after transfection. Lentiviral supernatant was collected 72 h after transfection and concentrated by ultracentrifugation at 24,000 rpm for 4 h at 4°C using a Beckman Coulter SW 32 Ti rotor.

### Generation of Cas9-expressing THP-1 cells and pooled CRISPR screen

THP-1 cells were transduced with LentiCas9-Blast (Addgene plasmid 52962) [42] and selected with blasticidin to generate a stable SpCas9-expressing population. Cas9 activity was confirmed using the pXPR-011 GFP reporter assay (Addgene plasmid 59702) [13]. Cas9-expressing THP-1 cells were then transduced with the pooled sgRNA lentiviral library cloned in at low multiplicity of infection. The average multiplicity of infection was 0.16, and a minimum representation of approximately 1,000 cells per sgRNA was maintained across three biological replicates. Twenty-four hours after transduction, cells were selected with blasticidin and puromycin to enrich for cells carrying both Cas9 and the sgRNA library. Cells were expanded for seven population doublings, corresponding to 14 days post-transduction.

### Transwell migration screen

The pooled CRISPR migration screen was performed using Corning Transwell inserts with 5 µm pore-size polycarbonate membranes. THP-1 cells were seeded into the upper chamber at 5 × 10⁵ cells/ml. The lower chamber contained medium supplemented with 200 ng/ml CCL2 as chemoattractant, with PBS used as a negative-control condition. After 12 h, cells were collected separately from the upper chamber, representing the non-migrant fraction, and the lower chamber, representing the migrant fraction. A minimum of 2.5 million cells was collected per condition per chamber. Three independent biological replicates were collected at separate time points. Plasmid library, day 14 pre-assay, migrant, and non-migrant samples were processed for sequencing.

### NGS library preparation and sequencing of sgRNA amplicons

Genomic DNA was extracted using the QIAGEN Blood & Tissue Kit. Integrated sgRNA cassettes were amplified from genomic DNA using LentiGuide-Puro-specific primers and Herculase II Fusion DNA Polymerase. For each sample, five 50 µl PCR reactions were performed using no more than 2.5 µg genomic DNA per reaction. Amplicons were purified by agarose gel extraction. A second PCR was performed with handle-specific indexing primers to add sequencing adapters and sample indexes using 0.5 µM forward and reverse primers. PCR products of the expected size were gel purified, quantified using the Roche Diagnostics KAPA Library Quantification Kit, catalog no. 50-196-5234, pooled, and sequenced on an Illumina NovaSeq X Plus platform using 150 bp paired-end reads.

### sgRNA extraction, counting, and quality control

Raw FASTQ files were used for sgRNA quantification and downstream CRISPR-screen analysis. Because the sgRNA sequence was contained in Read 1, R1 FASTQ files were processed to extract the 20 nt sgRNA sequence downstream of the LentiGuide-Puro anchor sequence. The anchor sequence 5′-GGAAAGGACGAAACACCG-3′ was used to identify sgRNA-containing reads. Reads lacking the anchor sequence were discarded, and exactly 20 nt downstream of the anchor were retained for sgRNA counting. sgRNA abundance was quantified against the cleaned sgRNA library using MAGeCK [31] count. Count tables generated by MAGeCK were used for library quality control, normalization, differential enrichment analysis, and visualization of migration and fitness phenotypes. Control sgRNAs, including non-targeting, neutral-region, and AAVS1 safe-harbor controls, were used for control-guide normalization. Library quality was assessed using mapping rate, zero-count sgRNA frequency, and Gini index.

### Differential sgRNA and gene-level enrichment analysis

Differential enrichment was assessed using MAGeCK *test*. The primary comparison contrasted migrated and non-migrated fractions from the transwell assay to identify sgRNAs associated with CCL2-directed migration. Raw sgRNA-level migration phenotypes were defined as log2 fold change in migrated cells relative to non-migrated cells. Negative values indicated depletion from the migrated fraction, whereas positive values indicated enrichment in the migrated fraction. Because cells carrying non-targeting or neutral-region control sgRNAs are expected to behave like unperturbed THP-1 cells, the control-guide distribution was used to define the neutral migration baseline. For control-centered analyses, the median migration LFC of all control sgRNAs was subtracted from each sgRNA migration LFC. This defined neutral migration behavior as zero and allowed gene-targeting sgRNAs to be interpreted relative to normally migrating control cells. Unless otherwise indicated, migration scores shown in Fig. 1D-H represent control-centered migration scores.

Gene-level migration effect sizes were calculated as the median control-centered sgRNA migration score for each gene. Gene-level significance was determined using MAGeCK Robust Rank Aggregation from the migrated versus non-migrated comparison. Genes were considered significant when the positive- or negative-selection FDR was <0.05. Genes depleted from the migrated fraction after control centering were interpreted as required for efficient CCL2-directed migration. Genes that remained near the control baseline were interpreted as not showing a reproducible migration phenotype under the conditions tested.

### Fitness-effect analysis

Fitness effects were estimated independently from the migrant versus non-migrant comparison using the MAGeCK-normalised count table. For each sgRNA, fitness log2 fold change was calculated as: fitness LFC = log2(mean day14 count + 1) − log2(plasmid count + 1). Gene-level fitness effects were summarized as the median sgRNA fitness log2 fold change per gene. These values were compared with gene-level migration scores to distinguish migration-specific phenotypes from general changes in sgRNA representation during culture. Fitness scores were used descriptively and were not assigned significance markers in the final visualizations. Control-guide behavior was assessed by comparing control sgRNA log2 fold-change distributions before and after control-guide normalization and median centering. Gene- level migration scores were displayed as horizontal bar plots, with colors indicating depletion, enrichment, or non-significance based on MAGeCK FDR <0.05. Gene-level migration and fitness scores were compared using scatter plots of median values.

### CRISPRtile analysis and residue-level score prediction

For structure-based analysis, sgRNA-level functional scores were merged with guide annotations and processed using the CRISPRtile [37] pipeline. Analyses were performed for selected WAVE regulatory complex subunits, including *WASF2*, *WASF3*, *ABI1*, *BRK1*, *CYFIP1*, and *NCKAP1L*. Input tables contained sgRNA sequences, genomic coordinates, gene annotations, and experimental functional scores. sgRNA columns were identified automatically by detecting valid 20 nt A/C/G/T guide sequences. Residue-level annotations were generated by mapping sgRNAs to coding sequences and integrating structural and sequence features. PDB-derived structural features were extracted using MDtraj v1.11.0 [35] and included DSSP secondary-structure annotations, solvent-accessible surface area calculated using the Shrake-Rupley [47] algorithm, and backbone atom coordinates. Protein-domain annotations were retrieved from InterPro, intrinsic disorder scores were obtained from IUPred, and PROVEAN scores [7] were used as a measure of evolutionary constraint. Guide-efficiency correction was performed using AutoGluon TabularPredictor [16] with default parameters on the guide dependent scores from CRISPOR [27]. Models were trained independently for each score type using the medium-quality preset and time limit. To estimate guide-efficiency-corrected residue-level effects, guide-specific features were imputed to their median or modal values before prediction. Corrected functional and dropout predictions were appended to the residue-level annotation table and used for downstream hotspot and structural analyses.

### Identification of migration-specific CYFIP1 hotspots

Migration-specific CYFIP1 hotspots were defined as residues with guide-efficiency-corrected Function scores ≤ −1.96 and Dropout scores > −0.25. These thresholds selected residues with strong migration-associated effects while excluding residues with strong fitness-associated dropout. Sensitivity of hotspot calls was evaluated across percentile-based cutoffs ranging from 10% to 30%.

### Structural mapping and contact analysis

The WAVE regulatory complex structure PDB 3P8C [6] was used as the primary structural reference. CYFIP1 scores were mapped directly to chain A. The active Rac1-bound WRC structure PDB 7USE [12] was used for comparison of Rac1-binding sites. For NCKAP1L visualization, NCKAP1 in the 3P8C structure was used as a structural proxy (NCKAP1(L). NCKAP1L and NCKAP1 sequences were aligned using UniProt Align to map equivalent residue positions. Inter-chain contacts between CYFIP1 and WRC partner proteins were defined as residue pairs containing at least one inter-chain heavy-atom distance ≤4 Å, calculated in PyMOL. CYFIP1 residues contacting more than one partner were assigned to a multiple-contact category. Contact annotations were exported as binary residue-level features and merged with CRISPRtile scores. Hotspot enrichment across structural contact categories was assessed using one-sided Mann-Whitney U tests, comparing each contact category with residues lacking annotated contacts. The directional hypothesis was that contact residues would have lower, more negative functional scores than non-contact residues. Pearson correlations were calculated using numpy. Amino-acid enrichment was calculated as the difference between hotspot amino-acid frequency and background amino-acid frequency within each contact group.

### Structural visualization

Structural figures were generated in PyMOL v2.x. Per-residue functional scores were mapped onto protein structures using score-based color gradients. Score gradients were clipped at the 5th and 95th percentiles to improve color contrast. Molecular surfaces were rendered using PyMOL surface representations with partial transparency. Final structural images were ray traced at 2,400 × 1,800 pixels.

Full-length protein visualizations were generated using AlphaFold-predicted structures obtained from the AlphaFold Protein Structure Database [28]

### ProTiler analysis of critical hypersensitive regions

ProTiler/TGUH was originally developed to identify CRISPR knockout hypersensitive regions from tiling-sgRNA dropout screens. Here, we adapted the TGUH segmentation strategy to guide-efficiency-corrected CRISPRtile Function scores to identify functionally hypersensitive regions associated with the migration phenotype. Guide-efficiency-corrected functional and dropout scores were extracted from the CRISPRtile prediction output and converted into ProTiler-compatible tab-separated format containing the required Symbol and AA columns. ProTiler was run separately on our CRISPRtile score for each analyzed WRC gene using the TGUH changepoint detection method. Functional and dropout score columns were specified as input score tracks. For genes that completed the full ProTiler workflow, critical hypersensitive-regions were identified from ProTiler output files. For CYFIP1, segmentation output was used directly to extract regions annotated as hypersensitive regions, and a custom matplotlib visualization was generated with solvent accessibility, DSSP secondary structure, and exon annotation tracks.

### Drug-target region selection and compound-interaction prediction

Candidate drug-target regions were selected from ProTiler-defined functionally hypersensitive regions using a composite scoring framework. The composite score incorporated four criteria: CRISPR-derived functional importance, defined by the mean TGUH segment Function score; target-length optimality relative to a 15-50 amino acid window; structural order based on DSSP secondary-structure annotation; and low intrinsic disorder based on IUPred. Criteria were weighted 50%, 20%, 15%, and 15%, respectively. Essentiality-related scores were normalized within each gene before composite scoring. Drug-protein interaction prediction was performed using BBBShinobi with the pretrained TransformerCPI2.0 model. A library of FDA-approved compounds was retrieved from ChEMBL and filtered for compounds with valid canonical SMILES strings. The NCKAP1L 91-121 amino acid sequence was used as the target peptide sequence. TransformerCPI2.0 scores were calculated for each compound-sequence pair using the BBBShinobi screening script. Computational single-residue deletion mutagenesis was performed for the selected compound-peptide pair. Each residue within the 30 amino acid target sequence was deleted individually, and the TransformerCPI2.0 interaction score was recalculated. Scores were logit transformed as log(score / [1 − score]) before downstream analysis.

### Drug-treatment transwell validation assay

For drug-validation experiments, THP-1 cells were pretreated overnight with Montelukast sodium, Piperacetazine, or DMSO vehicle control. Montelukast sodium was tested at 1 µM and 10 µM in wild-type and CYFIP1 knockout THP-1 cells. Piperacetazine was tested at 1 µM in wild-type THP-1 cells. Where indicated, wild-type and CYFIP1 knockout cells were analyzed in parallel.

Following drug pretreatment, cells were subjected to transwell migration assays using Corning Transwell inserts with 8 µm pores. Cells were seeded in the upper chamber at 5 x 10^5 cells/ml, and the lower chamber contained medium supplemented with 200 ng/ml CCL2 or PBS as a negative-control condition. After 6 h, cells from the upper and lower chambers were collected separately. Live and dead cell numbers were measured using a Cellometer instrument. Migration efficiency was calculated using live cells only:

% migration = bottom live cells / (top live cells + bottom live cells) × 100

Dead cells were excluded from the migration calculation to avoid confounding by differential viability. Viability was monitored independently in upper- and lower-chamber populations and calculated as:

% viability = live cells / (live cells + dead cells) × 100

For Montelukast validation, experiments were performed in three independent biological replicates per genotype and treatment condition unless otherwise indicated. Statistical analysis was performed using two-way ANOVA with genotype and treatment as factors, followed by pairwise comparisons where indicated.

For Piperacetazine validation, experiments were performed across two independent experiments with three replicates per condition per experiment, for a total of six replicates per condition. Piperacetazine-treated and vehicle-treated cells were compared using an unpaired two-tailed Student’s t-test. Data are presented as mean ± SEM.

### Bulk RNA-seq analysis of WRC gene expression

Previously generated bulk RNA-seq data from the same THP-1 cell line used in this study were re-analyzed to assess expression of WRC subunits and SPI1. Raw reads were pseudoaligned to the human GRCh38 transcriptome using kallisto. Transcript-level abundance estimates were aggregated to gene level and normalized to counts per million. Expression values were log2 transformed as log2(CPM + 1). Expression of WAVE regulatory complex subunits and the myeloid marker SPI1 was visualized across biological replicates. Heatmaps were generated in Python using seaborn, with gene expression values displayed as log2 CPM without z-score scaling.

### Human microglial single-nucleus RNA-seq analysis

Human brain cell-type expression patterns shown in Fig. 4A-B were examined using the single-nucleus RNA-seq dataset from the epigenomic dissection of Alzheimer’s disease study published in Cell [54]. Expression of WRC-associated genes was queried across major human brain cell types using the corresponding public single-cell portal, implemented through the UCSC Cell Browser framework described by Speir et al [49]. Expression of selected WRC genes, including *BRK1, WASF1, CYFIP1*, and *NCKAP1L*, was inspected across excitatory neurons, inhibitory neurons, oligodendrocytes, astrocytes, oligodendrocyte precursor cells, microglia, and vascular-associated cell populations. Portal-derived dot plots and feature plots were used to assess relative expression patterns across cell types. For independent validation of cell-type expression patterns in Fig. S4, WRC gene expression was queried in the public dataset from Green et al. [20]. Expression of selected WRC genes was inspected across major human brain cell types to validate microglia-enriched versus broadly expressed WRC components in an independent aging and Alzheimer’s disease cohort.

Microglial-state analyses shown in Fig. 4C-D were performed using the human microglial single-nucleus RNA-seq dataset from Sun et al. [51]. A cleaned microglial subset was loaded from an .h5ad file and analyzed using Scanpy. Expression matrices were inspected to confirm normalization status and, where required, counts were normalized to 10,000 counts per nucleus followed by log1p transformation. Microglial state annotations, Alzheimer’s disease diagnostic category, and brain-region metadata were preserved from the original dataset. Expression of WRC-associated genes, including *BRK1, CYFIP1, NCKAP1*, and *NCKAP1L*, was analyzed across microglial states and diagnostic categories. Genes not detected in the dataset were excluded from visualization. Dot plots were generated with dot size representing the fraction of nuclei expressing each gene and dot color representing scaled mean expression per gene across groups. Differential expression across Alzheimer’s disease diagnostic categories in the Sun et al. dataset was tested using Scanpy’s rank_genes_groups function with a Wilcoxon rank-sum test in a one-versus-rest design. Adjusted P values were calculated using the Benjamini-Hochberg method. Differential-expression results were extracted *for BRK1, CYFIP1, NCKAP1*, and *NCKAP1L*. Significance was defined as adjusted P < 0.05.

---

**Figure S1.**
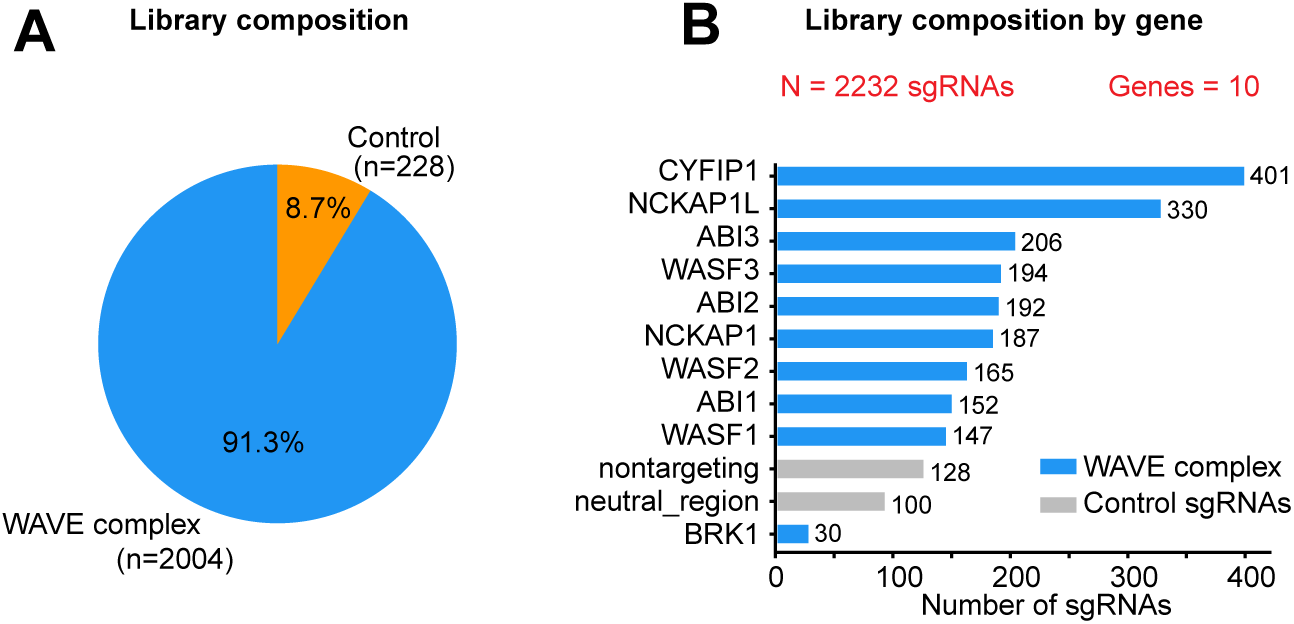
Composition of the WRC CRISPR tiling library. **A**. Proportion of sgRNAs targeting WRC subunits and control regions. **B**. Number of sgRNAs targeting each WRC subunit and control category. N, total number of sgRNAs in the library.

**Figure S2.**
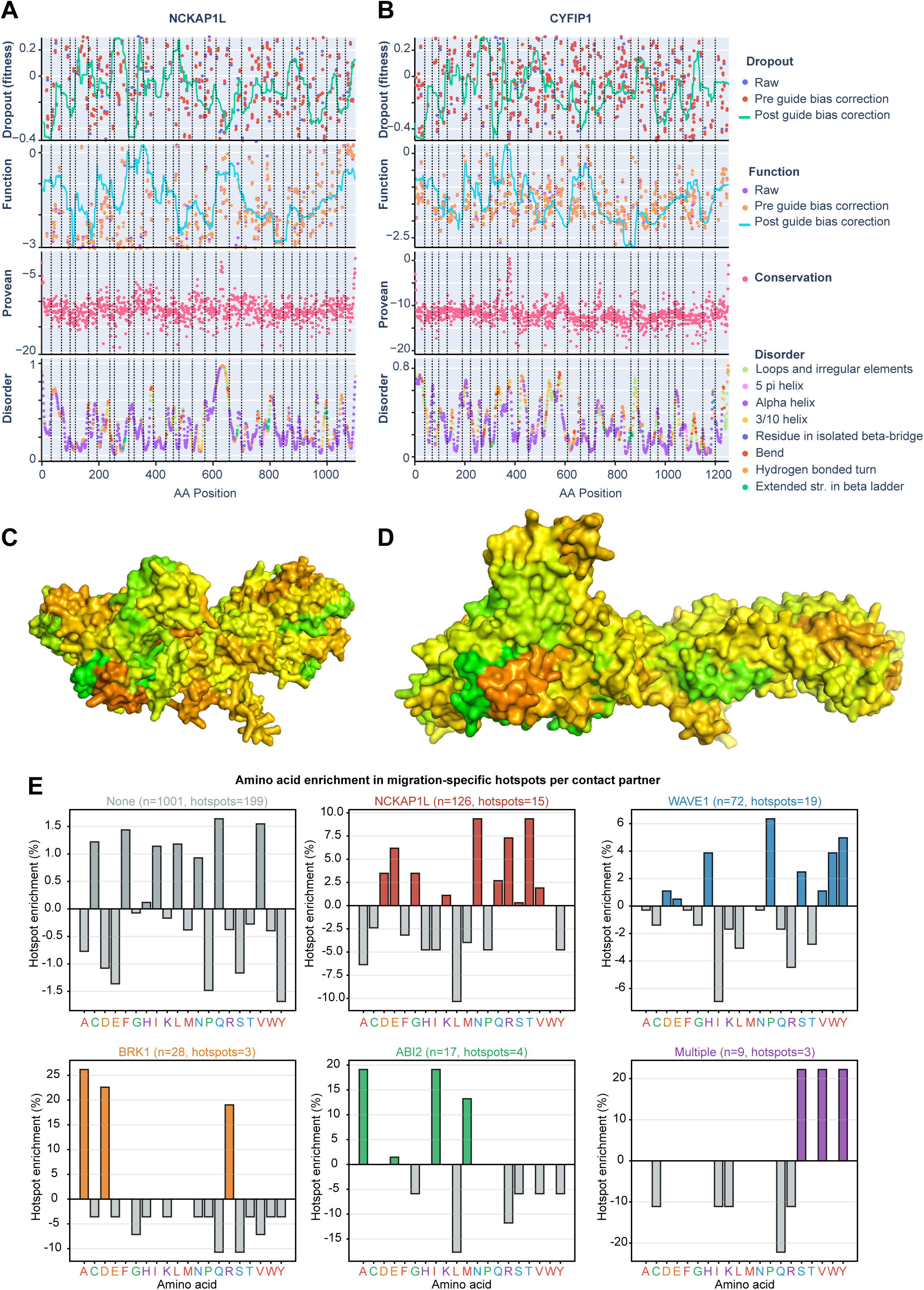
Functional, structural, and sequence analyses of NCKAP1L and CYFIP1. **A-B**. Per- residue CRISPRtile annotations for NCKAP1L (A) and CYFIP1 (B), including Function scores, Dropout scores, evolutionary conservation (PROVEAN), and predicted secondary structure/disorder. **C-D**. Mapping of Dropout scores onto AlphaFold structural models of *NCKAP1L* (C) and *CYFIP1* (D). **E**. Amino-acid enrichment among migration-specific hotspot residues grouped by WRC contact partner. Bars show the difference in amino-acid frequency between hotspot residues and the corresponding contact-group background composition.

**Figure S3.**
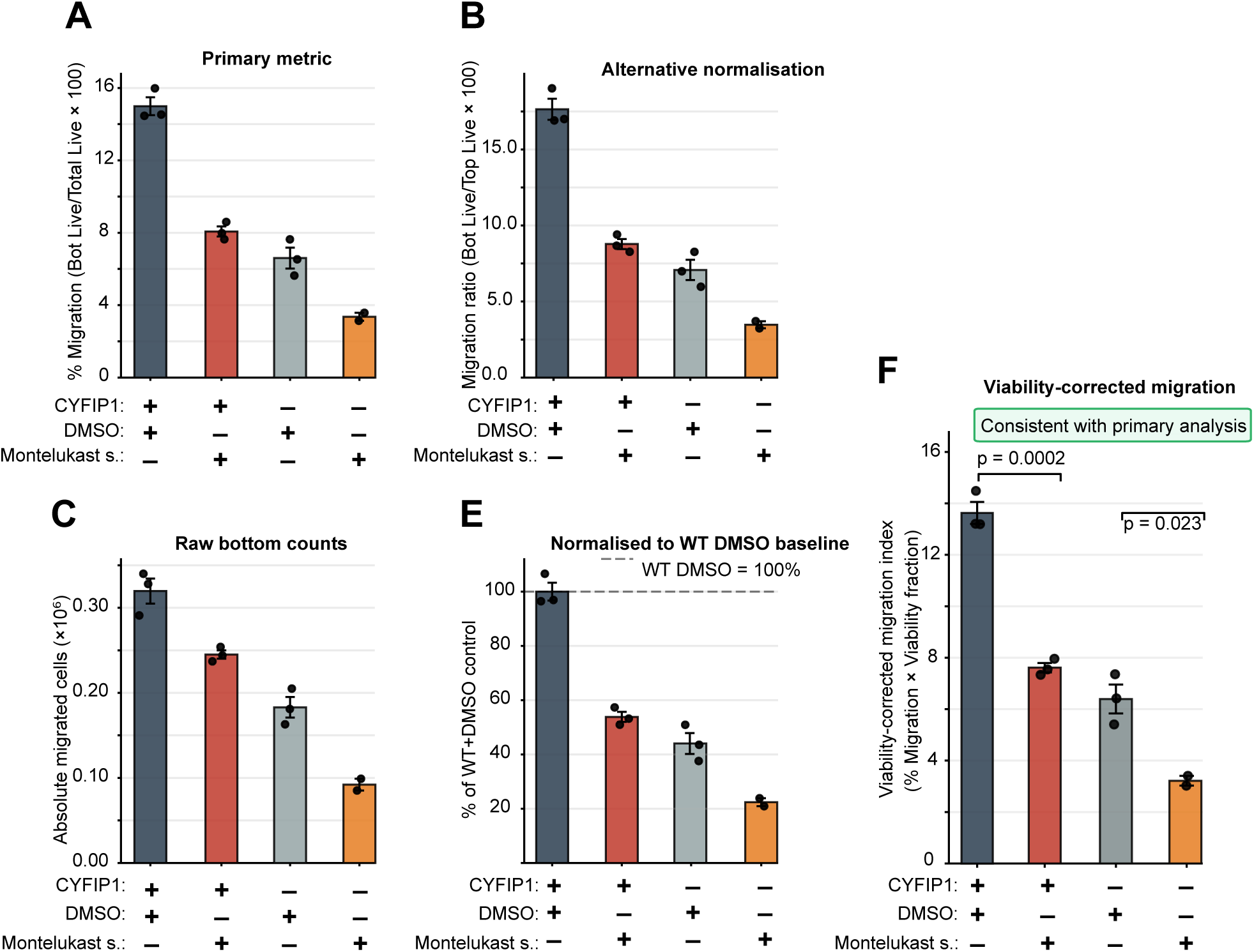
Analyses of Montelukast sodium effects on THP-1 migration. (A-F) Alternative migration quantification metrics, normalization approaches, and viability-corrected analyses supporting the primary migration phenotype shown in Fig. 3F.

**Figure S4.**
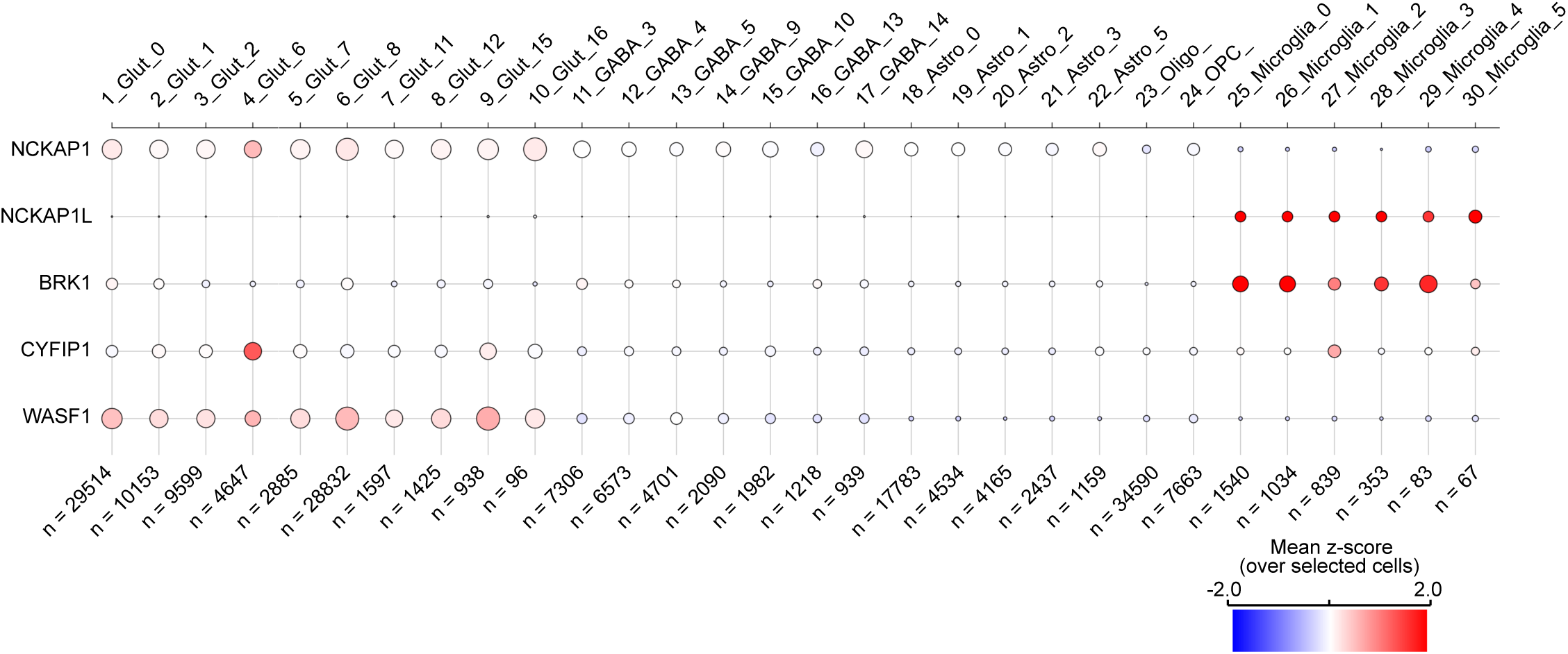
pendent validation of NCKAP1L microglial enrichment in human brain single- nucleus RNA-sequencing data. Expression of *NCKAP1, NCKAP1L, BRK1, CYFIP1*, and *WASF1* across neuronal, glial, and microglial populations from an independent human brain single-nucleus RNA-sequencing dataset [20]. Dot size indicates the number of cells within each cluster, and color represents mean normalized expression (z-score). *NCKAP1L* expression is largely restricted to microglial populations, whereas *CYFIP1* and *WASF1* exhibit broader expression across multiple cell types.

**Figure S5.**
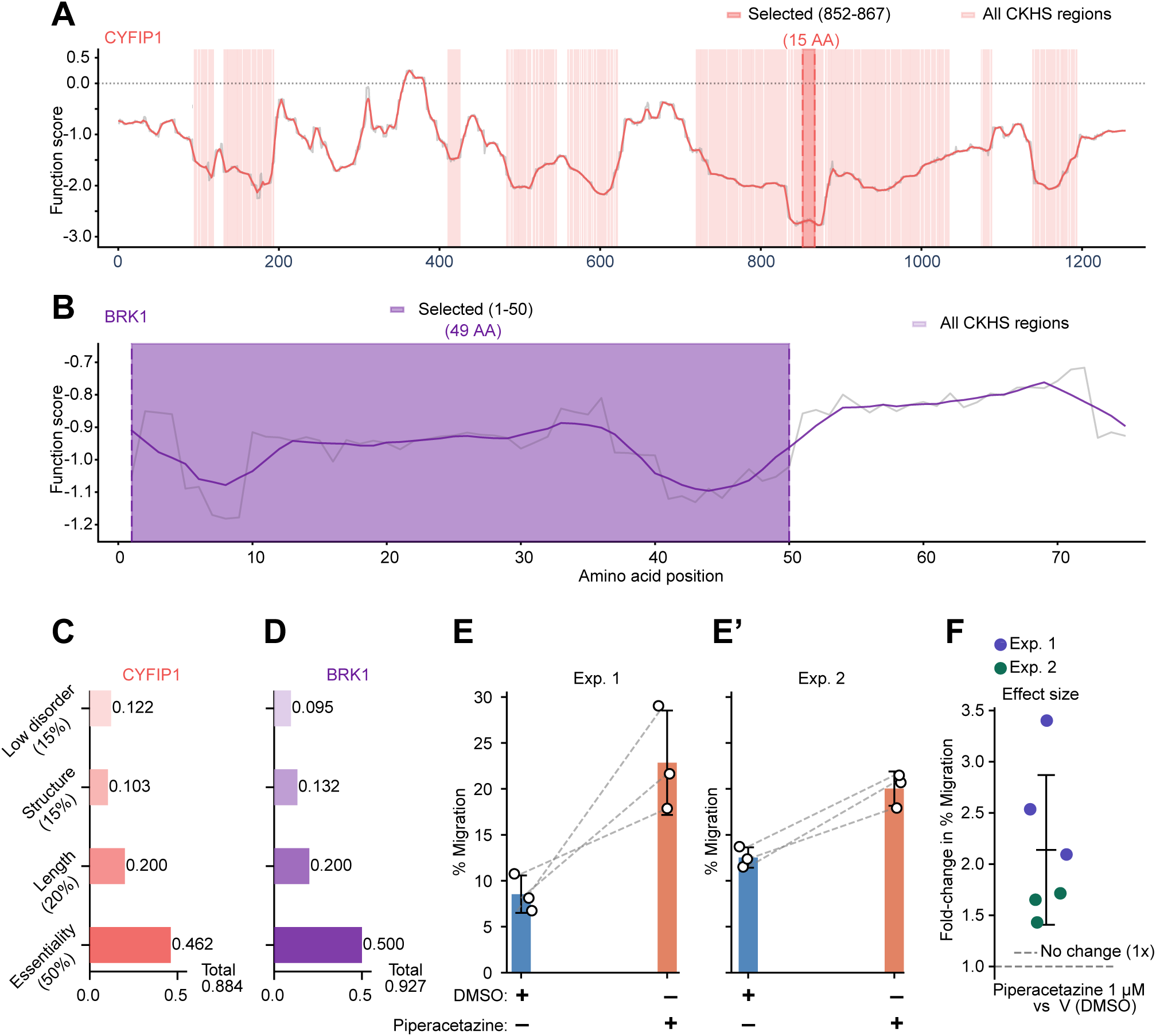
CRISPRtile target prioritization and Piperacetazine validation analyses. A-B. CRISPRtile Function score profiles and functionally hypersensitive region annotations for CYFIP1 (A) and BRK1 (B). Gray lines show guide-efficiency-corrected Function scores across each protein, and colored lines show 7-residue smoothed profiles. Shaded regions indicate functionally hypersensitive segments identified using TGUH segmentation, with the selected region for compound-interaction prediction highlighted. **C- D**. Composite prioritization of candidate drug-screening regions for CYFIP1 (C) and BRK1 (D). Horizontal bars show weighted contributions of functional importance, target length, structural order, and low intrinsic disorder to the final composite score for the selected region in each protein. **E-E′.** Individual transwell migration experiments showing the percentage of migrated THP-1 cells following treatment with vehicle or Piperacetazine (1 µM). Each point represents an individual replicate. **F**. Fold-change in CCL2-directed migration following Piperacetazine treatment relative to vehicle control. Each point represents an experimental replicate or experiment-level estimate, as indicated.

## References

1. (2025) 2025 Alzheimer’s disease facts and figures. Alzheimer’s & Dementia 21: e70235 Doi 10.1002/alz.70235

2 Bolmont T, Haiss F, Eicke D, Radde R, Mathis CA, Klunk WE, Kohsaka S, Jucker M, Calhoun ME (2008) Dynamics of the microglial/amyloid interaction indicate a role in plaque maintenance. J Neurosci 28: 4283–4292 Doi 10.1523/JNEUROSCI.4814-07.2008

3 Canver MC, Smith EC, Sher F, Pinello L, Sanjana NE, Shalem O, Chen DD, Schupp PG, Vinjamur DS, Garcia SP et al (2015) BCL11A enhancer dissection by Cas9-mediated in situ saturating mutagenesis. Nature 527: 192–197 Doi 10.1038/nature15521

4 Castro CN, Rosenzwajg M, Carapito R, Shahrooei M, Konantz M, Khan A, Miao Z, Gross M, Tranchant T, Radosavljevic M et al (2020) NCKAP1L defects lead to a novel syndrome combining immunodeficiency, lymphoproliferation, and hyperinflammation. J Exp Med 217: Doi 10.1084/jem.20192275

5 Chen L, Fan Z, Chang J, Yang R, Hou H, Guo H, Zhang Y, Yang T, Zhou C, Sui Q et al (2023) Sequence-based drug design as a concept in computational drug design. Nat Commun 14: 4217 Doi 10.1038/s41467-023-39856-w

6 Chen Z, Borek D, Padrick SB, Gomez TS, Metlagel Z, Ismail AM, Umetani J, Billadeau DD, Otwinowski Z, Rosen MK (2010) Structure and control of the actin regulatory WAVE complex. Nature 468: 533–538 Doi 10.1038/nature09623

7 Choi Y, Sims GE, Murphy S, Miller JR, Chan AP (2012) Predicting the functional effect of amino acid substitutions and indels. PLoS One 7: e46688 Doi 10.1371/journal.pone.0046688

8 Cocos R, Popescu BO (2024) Scrutinizing neurodegenerative diseases: decoding the complex genetic architectures through a multi-omics lens. Hum Genomics 18: 141 Doi 10.1186/s40246-024-00704-7

9 Cook S, Lenardo MJ, Freeman AF (2022) HEM1 Actin Immunodysregulatory Disorder: Genotypes, Phenotypes, and Future Directions. J Clin Immunol 42: 1583–1592 Doi 10.1007/s10875-022-01327-0

10 Courade JP, Zetterberg H, Hoglinger GU, Dewachter I (2025) The evolving landscape of Alzheimer’s disease therapy: From Abeta to tau. Cell 188: 7337–7354 Doi 10.1016/j.cell.2025.11.033

11 d’Errico P, Ziegler-Waldkirch S, Aires V, Hoffmann P, Mezo C, Erny D, Monasor LS, Liebscher S, Ravi VM, Joseph K et al (2022) Microglia contribute to the propagation of Abeta into unaffected brain tissue. Nat Neurosci 25: 20–25 Doi 10.1038/s41593-021-00951-0

12 Ding B, Yang S, Schaks M, Liu Y, Brown AJ, Rottner K, Chowdhury S, Chen B (2022) Structures reveal a key mechanism of WAVE regulatory complex activation by Rac1 GTPase. Nat Commun 13: 5444 Doi 10.1038/s41467-022-33174-3

13 Doench JG, Hartenian E, Graham DB, Tothova Z, Hegde M, Smith I, Sullender M, Ebert BL, Xavier RJ, Root DE (2014) Rational design of highly active sgRNAs for CRISPR-Cas9-mediated gene inactivation. Nat Biotechnol 32: 1262–1267 Doi 10.1038/nbt.3026

14 Duara R, Barker W (2022) Heterogeneity in Alzheimer’s Disease Diagnosis and Progression Rates: Implications for Therapeutic Trials. Neurotherapeutics 19: 8–25 Doi 10.1007/s13311-022-01185-z

15 Efthymiou AG, Goate AM (2017) Late onset Alzheimer’s disease genetics implicates microglial pathways in disease risk. Mol Neurodegener 12: 43 Doi 10.1186/s13024-017-0184-x

16 Erickson N, Mueller J, Shirkov A, Zhang H, Larroy P, Li M, Smola A (2020) Autogluon-tabular: Robust and accurate automl for structured data. arXiv preprint arXiv:200306505:

17. Fregoso FE, van Eeuwen T, Simanov G, Rebowski G, Boczkowska M, Zimmet A, Gautreau AM, Dominguez R (2022) Molecular mechanism of Arp2/3 complex inhibition by Arpin. Nat Commun 13: 628 Doi 10.1038/s41467-022-28112-2

18 Fujikawa R, Tsuda M (2023) The Functions and Phenotypes of Microglia in Alzheimer’s Disease. Cells 12: Doi 10.3390/cells12081207

19 Gao C, Jiang J, Tan Y, Chen S (2023) Microglia in neurodegenerative diseases: mechanism and potential therapeutic targets. Signal Transduct Target Ther 8: 359 Doi 10.1038/s41392-023-01588-0

20 Green GS, Fujita M, Yang HS, Taga M, Cain A, McCabe C, Comandante-Lou N, White CC, Schmidtner AK, Zeng Let al (2024) Cellular communities reveal trajectories of brain ageing and Alzheimer’s disease. Nature 633: 634–645 Doi 10.1038/s41586-024-07871-6

21 Griciuc A, Tanzi RE (2021) The role of innate immune genes in Alzheimer’s disease. Curr Opin Neurol 34: 228–236 Doi 10.1097/WCO.0000000000000911

22 Grinde B, Engdahl B (2017) Prescription database analyses indicates that the asthma medicine montelukast might protect against dementia: a hypothesis to be verified. Immun Ageing 14: 20 Doi 10.1186/s12979-017-0102-7

23 Gulisano W, Maugeri D, Baltrons MA, Fa M, Amato A, Palmeri A, D’Adamio L, Grassi C, Devanand DP, Honig LS et al (2018) Role of Amyloid-beta and Tau Proteins in Alzheimer’s Disease: Confuting the Amyloid Cascade. J Alzheimers Dis 64: S611–S631 Doi 10.3233/JAD-179935

24 Haq I, Ngo JC, Roy N, Lee E, Choudhury MA, Soni RK, Teich AF, Mayeux RP, De Jager PL, He Y et al (2026) Alzheimer’s disease risk protein SorLA regulates ER homeostasis and lipid metabolism in human microglia, with conserved effects in neurons. Acta Neuropathol 151: Doi 10.1007/s00401-026-03002-9

25 He W, Zhang L, Villarreal OD, Fu R, Bedford E, Dou J, Patel AY, Bedford MT, Shi X, Chen T et al (2019) De novo identification of essential protein domains from CRISPR-Cas9 tiling-sgRNA knockout screens. Nat Commun 10: 4541 Doi 10.1038/s41467-019-12489-8

26 Hsu JY, Fulco CP, Cole MA, Canver MC, Pellin D, Sher F, Farouni R, Clement K, Guo JA, Biasco L et al (2018) CRISPR-SURF: discovering regulatory elements by deconvolution of CRISPR tiling screen data. Nat Methods 15: 992–993 Doi 10.1038/s41592-018-0225-6

27 Hsu PD, Scott DA, Weinstein JA, Ran FA, Konermann S, Agarwala V, Li Y, Fine EJ, Wu X, Shalem O et al (2013) DNA targeting specificity of RNA-guided Cas9 nucleases. Nat Biotechnol 31: 827–832 Doi 10.1038/nbt.2647

28 Jumper J, Evans R, Pritzel A, Green T, Figurnov M, Ronneberger O, Tunyasuvunakool K, Bates R, Zidek A, Potapenko A et al (2021) Highly accurate protein structure prediction with AlphaFold. Nature 596: 583–589 Doi 10.1038/s41586-021-03819-2

29. Karahan H, Smith DC, Kim B, Dabin LC, Al-Amin MM, Wijeratne HRS, Pennington T, Viana di Prisco G, McCord B, Lin PB et al (2021) Deletion of Abi3 gene locus exacerbates neuropathological features of Alzheimer’s disease in a mouse model of Abeta amyloidosis. Sci Adv 7: eabe3954 Doi 10.1126/sciadv.abe3954

30 Kosoy R, Fullard JF, Zeng B, Bendl J, Dong P, Rahman S, Kleopoulos SP, Shao Z, Girdhar K, Humphrey J et al (2022) Genetics of the human microglia regulome refines Alzheimer’s disease risk loci. Nat Genet 54: 1145–1154 Doi 10.1038/s41588-022-01149-1

31 Li W, Xu H, Xiao T, Cong L, Love MI, Zhang F, Irizarry RA, Liu JS, Brown M, Liu XS (2014) MAGeCK enables robust identification of essential genes from genome-scale CRISPR/Cas9 knockout screens. Genome Biol 15: 554 Doi 10.1186/s13059-014-0554-4

32 Liang X, Wu H, Colt M, Guo X, Pluimer B, Zeng J, Dong S, Zhao Z (2021) Microglia and its Genetics in Alzheimer’s Disease. Curr Alzheimer Res 18: 676–688 Doi 10.2174/1567205018666211105140732

33 Lish AM, Young-Pearse TL (2026) Decoding Alzheimer’s genetic risk through intercellular communication in the human brain: Lessons from Clusterin. Curr Opin Neurobiol 97: 103165 Doi 10.1016/j.conb.2026.103165

34 Marschallinger J, Schaffner I, Klein B, Gelfert R, Rivera FJ, Illes S, Grassner L, Janssen M, Rotheneichner P, Schmuckermair C et al (2015) Structural and functional rejuvenation of the aged brain by an approved anti-asthmatic drug. Nat Commun 6: 8466 Doi 10.1038/ncomms9466

35 McGibbon RT, Beauchamp KA, Harrigan MP, Klein C, Swails JM, Hernandez CX, Schwantes CR, Wang LP, Lane TJ, Pande VS (2015) MDTraj: A Modern Open Library for the Analysis of Molecular Dynamics Trajectories. Biophys J 109: 1528–1532 Doi 10.1016/j.bpj.2015.08.015

36 McQuade A, Blurton-Jones M (2019) Microglia in Alzheimer’s Disease: Exploring How Genetics and Phenotype Influence Risk. J Mol Biol 431: 1805–1817 Doi 10.1016/j.jmb.2019.01.045

37 Ngo JC, Schoonenberg VAC, Nandakumar R, Wu X, Sher F (2026) AI platform for CRISPR functional mapping and function-based drug design. bioRxiv: Doi 10.64898/2026.05.06.722817

38 Pathania M, Davenport EC, Muir J, Sheehan DF, Lopez-Domenech G, Kittler JT (2014) The autism and schizophrenia associated gene CYFIP1 is critical for the maintenance of dendritic complexity and the stabilization of mature spines. Transl Psychiatry 4: e374 Doi 10.1038/tp.2014.16

39 Rey-Suarez I, Wheatley BA, Koo P, Bhanja A, Shu Z, Mochrie S, Song W, Shroff H, Upadhyaya A (2020) WASP family proteins regulate the mobility of the B cell receptor during signaling activation. Nat Commun 11: 439 Doi 10.1038/s41467-020-14335-8

40 Rotty JD, Wu C, Bear JE (2013) New insights into the regulation and cellular functions of the ARP2/3 complex. Nat Rev Mol Cell Biol 14: 7–12 Doi 10.1038/nrm3492

41. Roy N, Haq I, Ngo JC, Bennett DA, Teich AF, De Jager PL, Olah M, Sher F (2024) Elevated expression of the retrotransposon LINE-1 drives Alzheimer’s disease-associated microglial dysfunction. Acta Neuropathol 148: 75 Doi 10.1007/s00401-024-02835-6

42 Sanjana NE, Shalem O, Zhang F (2014) Improved vectors and genome-wide libraries for CRISPR screening. Nat Methods 11: 783–784 Doi 10.1038/nmeth.3047

43 Schoonenberg VAC, Cole MA, Yao Q, Macias-Trevino C, Sher F, Schupp PG, Canver MC, Maeda T, Pinello L, Bauer DE (2018) CRISPRO: identification of functional protein coding sequences based on genome editing dense mutagenesis. Genome Biol 19: 169 Doi 10.1186/s13059-018-1563-5

44 Schwimmbeck F, Staffen W, Hohn C, Rossini F, Renz N, Lobendanz M, Reichenpfader P, Iglseder B, Aigner L, Trinka E et al (2021) Cognitive Effects of Montelukast: A Pharmaco-EEG Study. Brain Sci 11: Doi 10.3390/brainsci11050547

45 Scott-Solache J, Pei J, Drew J, Lopez-Domenech G, Jolivet RB, Nieto-Rostro M, Davenport EC, Arancibia-Carcamo IL, Attwell D, Kittler JT (2026) Control of microglial dynamics by the Arp2/3 complex and the autism- and schizophrenia-associated protein CYFIP1. Proc Natl Acad Sci U S A 123: e2532488123 Doi 10.1073/pnas.2532488123

46 Sher F, Hossain M, Seruggia D, Schoonenberg VAC, Yao Q, Cifani P, Dassama LMK, Cole MA, Ren C, Vinjamur DS et al (2019) Rational targeting of a NuRD subcomplex guided by comprehensive in situ mutagenesis. Nat Genet 51: 1149–1159 Doi 10.1038/s41588-019-0453-4

47 Shrake A, Rupley JA (1973) Environment and exposure to solvent of protein atoms. Lysozyme and insulin. J Mol Biol 79: 351–371 Doi 10.1016/0022-2836(73)90011-9

48. Sims R, van der Lee SJ, Naj AC, Bellenguez C, Badarinarayan N, Jakobsdottir J, Kunkle BW, Boland A, Raybould R, Bis JC et al (2017) Rare coding variants in PLCG2, ABI3, and TREM2 implicate microglial-mediated innate immunity in Alzheimer’s disease. Nat Genet 49: 1373–1384 Doi 10.1038/ng.3916

49 Speir ML, Bhaduri A, Markov NS, Moreno P, Nowakowski TJ, Papatheodorou I, Pollen AA, Raney BJ, Seninge L, Kent WJ et al (2021) UCSC Cell Browser: visualize your single-cell data. Bioinformatics 37: 4578–4580 Doi 10.1093/bioinformatics/btab503

50 Spencer WJ, Schneider NF, Lewis TR, Castillo CM, Skiba NP, Arshavsky VY (2023) The WAVE complex drives the morphogenesis of the photoreceptor outer segment cilium. Proc Natl Acad Sci U S A 120: e2215011120 Doi 10.1073/pnas.2215011120

51 Sun N, Victor MB, Park YP, Xiong X, Scannail AN, Leary N, Prosper S, Viswanathan S, Luna X, Boix CA et al (2023) Human microglial state dynamics in Alzheimer’s disease progression. Cell 186: 4386–4403 e4329 Doi 10.1016/j.cell.2023.08.037

52 Twiss E, McPherson C, Weaver DF (2025) Global Diseases Deserve Global Solutions: Alzheimer’s Disease. Neurol Int 17: Doi 10.3390/neurolint17060092

53 Valiukas Z, Tangalakis K, Apostolopoulos V, Feehan J (2025) Microglial activation states and their implications for Alzheimer’s Disease. J Prev Alzheimers Dis 12: 100013 Doi 10.1016/j.tjpad.2024.100013

54 Xiong X, James BT, Boix CA, Park YP, Galani K, Victor MB, Sun N, Hou L, Ho LL, Mantero J et al (2023) Epigenomic dissection of Alzheimer’s disease pinpoints causal variants and reveals epigenome erosion. Cell 186: 4422–4437 e4421 Doi 10.1016/j.cell.2023.08.040

55 Yamamoto A, Suzuki T, Sakaki Y (2001) Isolation of hNap1BP which interacts with human Nap1 (NCKAP1) whose expression is down-regulated in Alzheimer’s disease. Gene 271: 159–169 Doi 10.1016/s0378-1119(01)00521-2

56 Yates AG, Kislitsyna E, Alfonso Martin C, Zhang J, Sewell AL, Goikolea-Vives A, Cai V, Alkhader LF, Skaland A, Hammond B et al (2022) Montelukast reduces grey matter abnormalities and functional deficits in a mouse model of inflammation-induced encephalopathy of prematurity. J Neuroinflammation 19: 265 Doi 10.1186/s12974-022-02625-5

57 Young-Pearse TL, Lee H, Hsieh YC, Chou V, Selkoe DJ (2023) Moving beyond amyloid and tau to capture the biological heterogeneity of Alzheimer’s disease. Trends Neurosci 46: 426–444 Doi 10.1016/j.tins.2023.03.005

58 Zhang Y, Chen H, Li R, Sterling K, Song W (2023) Amyloid beta-based therapy for Alzheimer’s disease: challenges, successes and future. Signal Transduct Target Ther 8: 248 Doi 10.1038/s41392-023-01484-7

59 Zhao W, Liu Z, Wu J, Liu A, Yan J (2026) Potential targets of microglia in the treatment of neurodegenerative diseases: Mechanism and therapeutic implications. Neural Regen Res 21: 1497–1511 Doi 10.4103/NRR.NRR-D-24-01343

